# OsTIR1^F74G^ Expression Controls Basal and Auxin-Induced Degradation Rates of the AID System in Yeast

**DOI:** 10.1101/2025.07.11.664329

**Authors:** Julius A. Fülleborn, Andrei Seica, Andreas Milias-Argeitis, Matthias Heinemann

## Abstract

The auxin-inducible degron (AID) system has been widely used to conditionally and dynamically deplete proteins in yeast. In this system, the plant hormone auxin promotes OsTIR1-mediated degradation of proteins carrying an auxin-inducible degron tag. However, “basal” degradation of AID-tagged proteins in the absence of auxin has hampered work with essential proteins due to defective or non-viable strains. A second-generation AID system based on the OsTIR^F47G^ mutant was recently introduced to overcome the limitations of basal degradation in budding yeast. However, it remained unclear to what extent the use of OsTIR^F47G^ eliminates basal degradation and how it impacts auxin-induced degradation. Here, by performing a quantitative characterization of the basal and auxin-induced degradation dynamics using OsTIR1^F74G^ in budding yeast, we find that basal degradation is still detectable and that it depends on OsTIR1^F74G^ expression levels. We show that also the auxin-induced degradation rates and auxin-induced steady-state concentrations of AID-tagged proteins depend on OsTIR1^F74G^ expression levels, in addition to the type of auxin used. Lastly, we showcase how increased basal degradation of AID-tagged functional proteins can impair cell growth and lead to unwanted phenotypes. We anticipate that our findings will guide future applications of the AID system in budding yeast by helping researchers select appropriate OsTIR1^F74G^ expression levels, particularly in studies focused on precise control of degradation dynamics.

## Introduction

The auxin-inducible degron (AID) system has been widely used to dynamically deplete proteins in yeasts and other organisms^1–7^ – a crucial ability to study dynamic processes^8^. Auxin-induced degradation of AID-tagged proteins enables the study of dynamic adaptation of cells upon degradation of specific proteins^9–11^ as well as loss-of-function phenotypes of essential proteins, which otherwise cannot be deleted. For instance, a genome-wide library of protein-AID fusions has recently been developed enabling conditional investigation of essential genes^12^, mimicking the classical, temperature-sensitive yeast strains. Thus, the auxin-induced degradation of AID-tagged proteins is a powerful research tool.

The AID-system requires the interaction of three components: (i) the plant hormone auxin, (ii) the heterologously expressed plant F-box protein Transport Inhibitor Response 1 (TIR1), typically derived from *Oryza sativa* (OsTIR1)^1,3,13^, and (iii) a target protein carrying a specific, auxin-inducible degron tag (AID-tag)^1–3^. Auxin binds to TIR1 and stabilises the interaction between the AID-tag and the TIR1 binding pocket, promoting the SCF ubiquitin-ligase-mediated degradation of AID-tagged proteins^14,15^.

A drawback of the original AID system is that auxin-independent activity of OsTIR1 can cause unwanted degradation of AID-tagged proteins, resulting in problems with strain viability and/or the function of AID-tagged proteins^1–3,16–18^. This “basal” degradation of AID-tagged proteins can hamper studies where the precise abundance of the protein, targeted for auxin-induced degradation, is critical^17,19–22^. The problem of basal degradation was so far tackled by either structural modifications of the OsTIR1 auxin-binding pocket^2,21^, inducible expression of OsTIR1^17,18^ or usage of competitive inhibitors of OsTIR1^16^. Recently, a second-generation AID system was proposed^2^ which addressed the problem of basal degradation by engineering a modified OsTIR1^F74G^ version to specifically recognise the auxin analogue 5-Ph-IAA and thereby minimise unspecific interactions with non-auxin molecules in the culture media that might induce basal degradation. While initial findings with OsTIR1^F74G^ in budding yeast (*Saccharomyces cerevisiae*) suggested absence of basal degradation^2^, it remains unclear to which extent the use of OsTIR1^F74G^ fully eliminates basal degradation and impacts auxin-induced degradation in budding yeast.

Here, by expressing OsTIR1^F74G^ from six promoters with a wide range of expression strengths, we find that basal degradation still occurs when using OsTIR1^F74G^ and that it depends on the OsTIR1^F74G^ expression levels used. Furthermore, also auxin-induced degradation rates and auxin-induced steady state concentrations of AID-tagged proteins depend on the expression level of OsTIR1^F74G^ and the type of auxin used. Lastly, we demonstrate that OsTIR1^F74G^-mediated basal degradation of the essential protein Acc1 can cause growth defects and basal degradation of Tup1 can lead to flocculation. Our findings will help researchers choose appropriate OsTIR1^F74G^ expression strengths in yeast to balance the required degradation dynamics with a tolerable degree of basal degradation.

## Results

### Experimental design to quantify basal- and auxin-induced degradation by OsTIR1^F74G^

To assess and quantify basal degradation and auxin-induced degradation of AID-tagged proteins, we constructed a set of six budding yeast strains expressing genomically encoded, yeast-codon optimized OsTIR1^F74G^ from constitutive promoters with different strengths (Supplementary Figure 1)^23–25^. These strains also express a monomeric enhanced GFP^26^ C-terminally tagged with the short AID* sequence^22^ (amino acids 71–114 from AtIAA17, Uniprot ID: P93830, in the following referred to as AID) (**Figure 1**). We considered the single-cell GFP-AID fluorescence intensity as proxy for cellular GFP-AID concentration. By measuring the GFP-AID fluorescence intensity in each strain with a flow cytometer, we assessed the dependence of basal and auxin induced GFP-AID degradation on OsTIR1^F74G^ expression levels (**Figure 1**).

**Figure 1:**
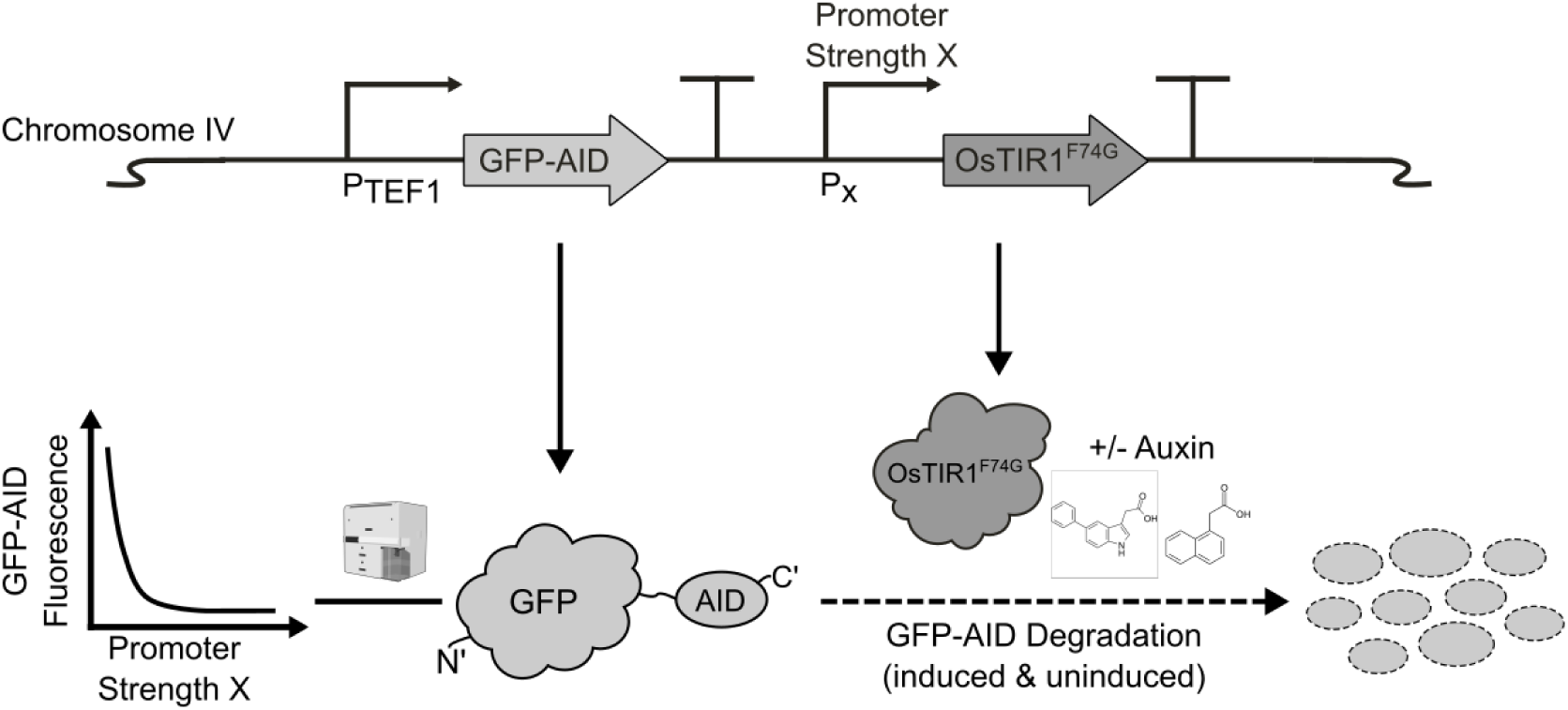
Experimental system to measure basal and auxin-induced degradation by the AID system in *S. cerevisiae*.

### Basal degradation is inversely correlated with OsTIR1^F74G^ expression strength

To test whether basal degradation is dependent on OsTIR1^F74G^ expression strength, we measured GFP-AID fluorescence in exponential phase cultures of each of the six strains using flow cytometry. First, we divided the GFP-fluorescence value of each single cell by its forward-scatter, a proxy for cell size^27^, to obtain an estimate of the GFP concentration in the cell and then took the median of the single cell GFP concentration estimates. Second, we corrected these median GFP concentration values for cellular background auto-fluorescence by subtracting the median auto-fluorescence of the wild-type cells from each median GFP concentration value. Then, we normalised the background-corrected GFP concentrations to the background-corrected GFP concentration of a control strain expressing GFP-AID but not OsTIR1^F74G^. Plotting the normalized GFP values against the strength of the promoters expressing OsTIR1^F74G^, we observed that even with the weakest promoter, the GFP fluorescence was ∼4% lower than in the strain without OsTIR1^F74G^ and found an anti-correlation between the GFP-AID fluorescence and the promoter strengths (**Figure 2**). These findings demonstrate that basal degradation indeed occurs with the AID system in yeast and suggest that the expression level of OsTIR1^F74G^ influences the degree of basal degradation.

**Figure 2:**
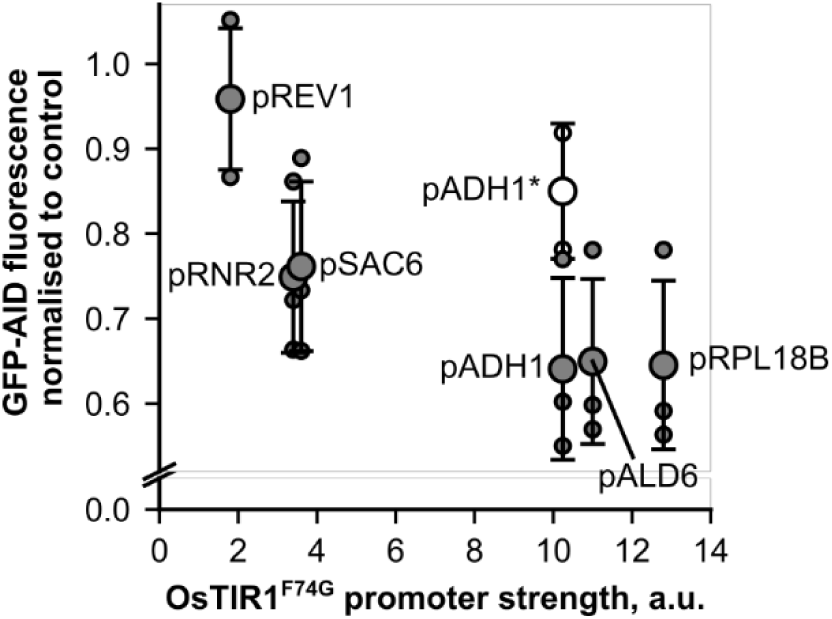
Basal degradation is inversely correlated with OsTIR1^F74G^ expression strength. GFP-AID fluorescence measurements in absence of auxin plotted against the promoter strength of each promoter controlling OsTIR1^F74G^ expression. GFP-AID fluorescence measurements are normalized to a control strain which expresses GFP-AID, but no OsTIR1^F74G^. Filled symbols represent codon-optimized OsTIR1^F74G^-genes while empty symbols denote non-codon-optimized OsTIR1^F74G^-genes. Small symbols represent values for individual biological replicates, whereas large symbols indicate the mean value of those values. Per biological replicate, five technical replicates were taken to account for measurement variability. Error-bars indicate the standard deviation across the biological replicates. Promoter strengths of all promoters except the ADH1 promoter are taken from Lee *et al.*^23^ based on the mRuby2 fluorescence measurements presented in Figure 3 of the same publication. We estimated the relative ADH1 promoter strength based on published data of protein abundance measurements of the native gene products expressed from the remaining promoters^25^ (for details refer to **Supplementary Figure 1**).

To verify that OsTIR1^F74G^ expression level is a determinant of the basal degradation rate, we constructed another strain expressing non-codon optimized OsTIR1^F74G^ from the strong ADH1 promoter^1,24^, which we expected would lead to lower a OsTIR1^F74G^ concentration compared to the codon-optimized OsTIR1^F74G^ used in our previous experiments. In line with our expectation, this strain indeed exhibited lower background degradation than its counterpart with codon-optimised OsTIR1^F74G^ (**Figure 2**). This finding provides additional support for the conclusion that the expression level of OsTIR1^F74G^ determines the degree of basal degradation.

### Auxin-induced degradation dynamics and steady state degradation depend on OsTIR1^F74G^ expression levels and the choice of auxin

Next, we asked how the different levels of OsTIR1^F74G^ expression would influence the dynamics of auxin-induced GFP-AID degradation. To assess the dynamics of auxin-induced degradation, we performed time-course measurements of the single-cell GFP-AID fluorescence signals after addition of 5-Ph-IAA using flow cytometry. For each strain, we fitted the dynamic data after auxin addition to a one-phase exponential decay function (**Figure 3A**; see **Supplementary Figure 2A** for all individual fits per strain). Thereby, for each strain, we obtained decay functions with the respective parameters describing the auxin-induced degradation dynamics of GFP-AID (**Figure 3B**). Here, we found that the degradation rates increase with the promoter strengths, except for the strain expressing non-codon optimised OsTIR1^F74G^ from pADH1 (**Figure 3C**). We found a ∼5-fold difference between the highest and lowest degradation rate in the strains tested. Using an alternative measurement approach, in which we continuously recorded GFP-AID fluorescence for 30 minutes after addition of 5-Ph-IAA with a high measurement frequency using flow cytometry (**Supplementary Figure 3A**), and fitting the same model to this alternative measurement data, we obtained similar degradation rates (**Supplementary Figure 3B**), lending further support to the degradation rates reported here. Increasing the concentration of 5-Ph-IAA in the medium did not increase the degradation rate of the strain expressing OsTIR1^F74G^, when we tested the promoter with the lowest expression strength (REV1 promoter) (**Supplementary Figure 4A, B**), which suggested that the OsTIR1^F74G^ degradation system is already saturated at the auxin concentrations applied. Overall, these findings demonstrate that the OsTIR1^F74G^ expression level not only determines the degree of basal degradation in the absence of auxin, but also the auxin-induced degradation rates.

**Figure 3:**
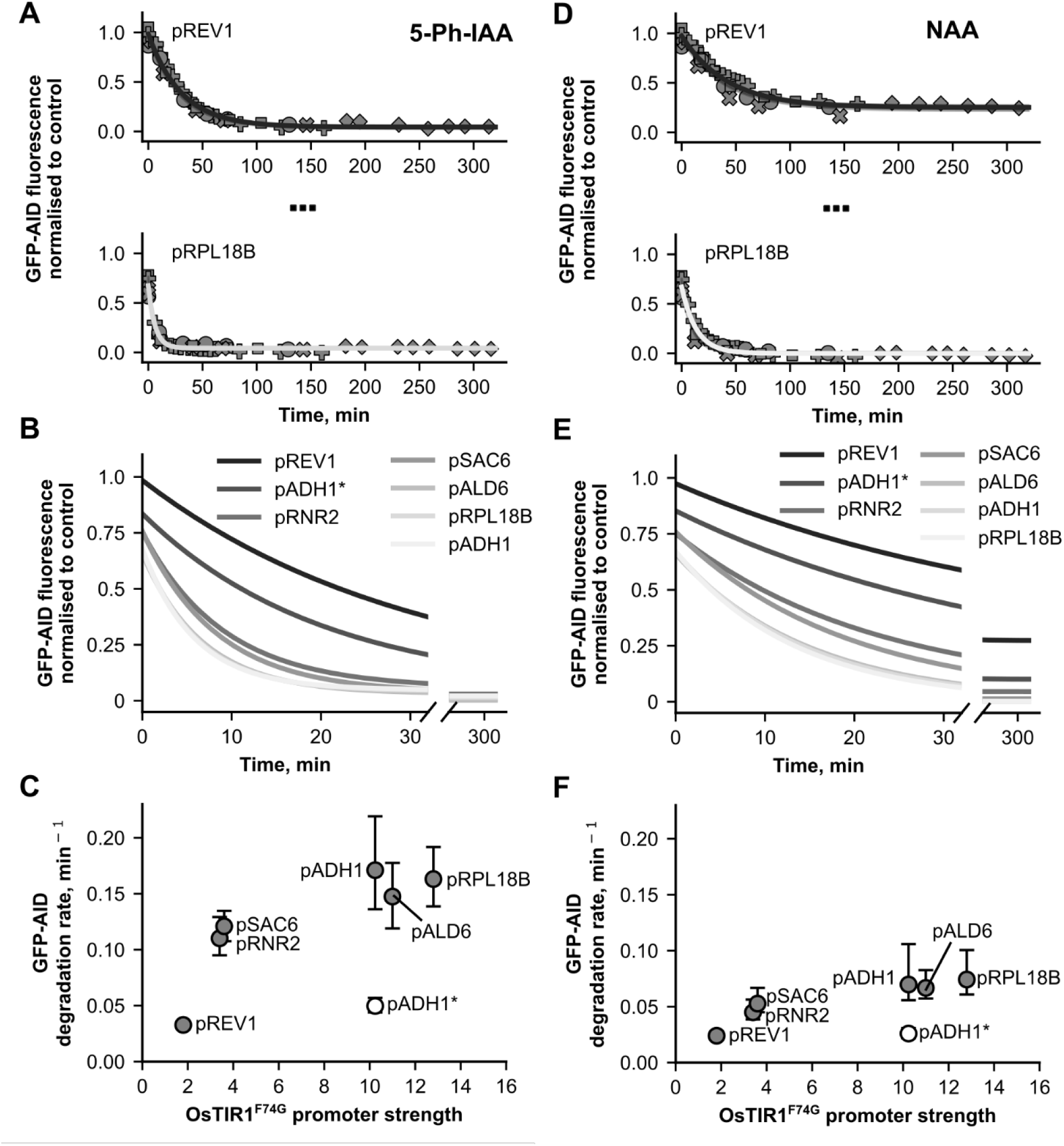
Auxin induced degradation dynamics depend on OsTIR1^F74G^ expression levels and the choice of auxin. **A, D** The GFP-AID fluorescence measured in the absence of auxin (timepoint 0) and in time-intervals after induction of GFP-AID degradation with 5-Ph-IAA (**A)** or NAA (**D**) using flow-cytometry. GFP-AID fluorescence measurements are normalized to a control strain which expresses GFP-AID, but no OsTIR1^F74G^. Each datapoint and each symbol-type corresponds to the median, normalized GFP-fluorescence measured per biological replicate at a given timepoint after auxin addition. The solid lines describe the fit of a one-step exponential decay function to the time course data. For readability, only the datapoints and fits of the strains expressing OsTIR1^F74G^ from the weakest and strongest promoter are shown, respectively. For a complete overview of data points and fits, refer to **Supplementary Figure 3**. **C, F** Overview of GFP-AID degradation rates derived from the fits depicted in **B** and **E**. Filled data points denote strains expressing codon optimized OsTIR1^F74G^, while the empty symbol denotes a strain expressing non-codon optimized OsTIR1^F74G^ from the strong ADH1 promoter (denoted by pADH1* on the graph). Error bars indicate the 95% confidence interval for the estimates of the degradation rates. For an overview of the degradation rates including uncertainty estimates, see **Supplementary Table 4** and **Supplementary Table 5**.

Since the OsTIR1^F74G^ variant used in this study was optimized to bind 5-Ph-IAA, we next asked if the degradation rates would be different if another type of auxin is used. We therefore performed the same time course experiments also with the synthetic auxin, 1-naphthalene acetic acid (NAA), which was previously found to have beneficial properties for fluorescence live cell imaging^10^. Here, we found a ∼3-fold difference between the highest and lowest degradation rate (**Figure 3D-F**; see **Supplementary Figure 2B** for all individual fits per strain). Degradation rates from NAA-induced and 5-Ph-IAA-induced degradation are linearly correlated (r=0.97, **Supplementary Figure 5**), where NAA-induced degradation rates are ∼56% lower than 5-Ph-IAA-induced degradation rates. Taken together, these findings demonstrate that auxin induced degradation rates with OsTIR1^F74G^ are faster when inducing degradation with the cognate auxin analogue 5-Ph-IAA than with the alternative auxin analogue NAA.

While so far, we focussed on the dynamics of degradation, we next investigated the steady-state GFP concentrations during continued auxin-exposure. Here, we found that the steady-state GFP-AID concentrations in all strains treated with 5-Ph-IAA were <10% of the GFP-AID concentrations in an untreated control strain, irrespective of OsTIR1^F74G^ promoter strength (**Figure 4A**). When GFP-AID depletion was instead induced with NAA, we found equally low steady state GFP-AID concentrations in most strains. The two exceptions, which displayed higher steady state GFP-AID concentrations, were the strain expressing the codon-optimized OsTIR1^F74G^ from the weak REV1 promoter and the strain expressing the non-codon-optimized OsTIR1^F74G^ from the strong ADH1 promoter (**Figure 4B**). These findings show that low steady state GFP-AID concentrations can only be reached when either 5-Ph-IAA is used or if NAA is used and OsTIR1^F74G^ is highly expressed suggesting that NAA is a less potent inducer of OsTIR1^F74G^ activity than 5-Ph-IAA.

**Figure 4:**
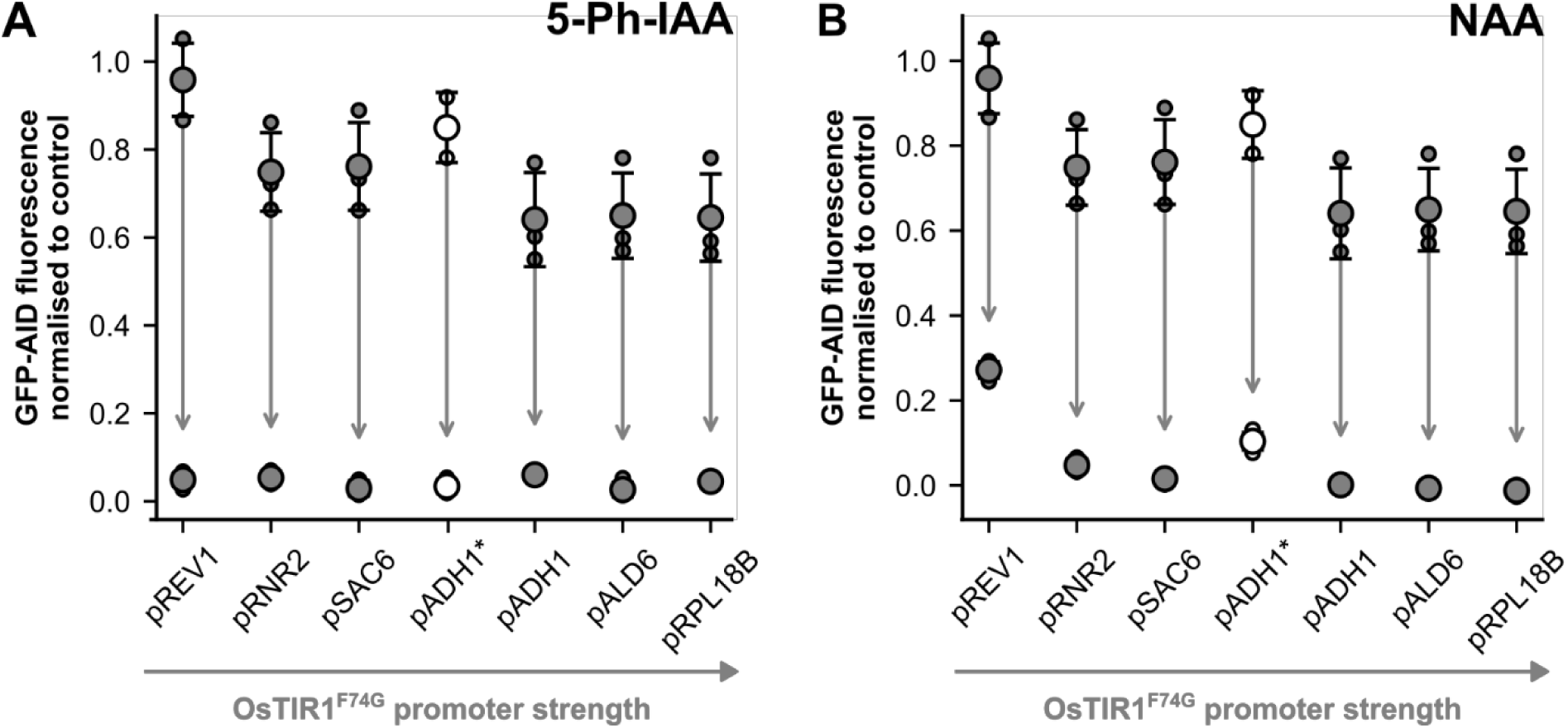
The steady state GFP-AID levels after auxin-induced degradation are dependent on OsTIR1^F74G^ expression and the choice of auxin. Fluorescence measurements of GFP-AID before addition of auxin and after addition of 5-Ph-IAA (**A**) or NAA (**B**) when GFP-AID levels reached a steady state. The directed arrows have their origins in GFP-AID measurements before addition of auxin and point towards the datapoints after reaching steady state levels in the presence of auxin. The promoters on the x-axis are sorted categorically based on their strength which is indicated by the grey, horizontal arrow. Small datapoints describe the median GFP-AID signal per biological replicate while big datapoints are the mean GFP-AID fluorescence across those biological replicates. Filled symbols represent strains expressing codon-optimized OsTIR1^F74G^, whereas open symbols represent non-codon-optimized OsTIR1^F74G^ which is also indicated by the * behind the promoter name. Error bars represent the standard deviation of the GFP-signal per biological replicate.

### High basal degradation of AID-tagged protein can lead to growth defects

Having characterised basal degradation in yeast expressing OsTIR1^F74G^, we next tested the effect of basal degradation on a strain’s phenotype when an essential protein is tagged with AID. As a target, we chose ACC1, an essential gene whose gene product catalyses the rate limiting step in fatty acid biosynthesis^28,29^. Due to its essential role in metabolism, we expected that high basal degradation of AID-tagged Acc1 would have an observable impact on the strain’s phenotype – such as the growth rate. To test this, we measured the growth rate of ACC1-AID yeast strains with expression of OsTIR1^F74G^ from either a weak (i.e. pREV1) or a strong promoter (i.e. pADH1). To control for possible growth impairment arising from OsTIR1^F74G^ expression or AID-tagged ACC1 alone, we also determined the growth rates of strains only expressing OsTIR1^F74G^ from the ADH1 or REV1 promoter and strains with ACC1-AID without OsTIR1^F74G^ (see **Supplementary Figure 6** for an overview of all individual growth curves).

Here, we found that the control strains expressing either OsTIR1^F74G^ or ACC1-AID individually had growth rates similar to the wildtype, suggesting that these genomic alterations alone were not leading to significant growth defects. In contrast, the strain expressing OsTIR1^F74G^ from the ADH1 promoter and having ACC1 AID-tagged had growth rates that were significantly lower than the wild type and the strain only carrying the ACC1-AID tag (**Figure 5**; Kruskal-Wallis test, Dunn’s pos hoc test with Benjamini-Hochberg correction, p-value < 0.05 *). These results demonstrate that basal degradation of certain proteins can have drastic growth phenotypes, highlighting the importance of choosing the right OsTIR1^F74G^ expression levels, when tagging essential proteins.

**Figure 5:**
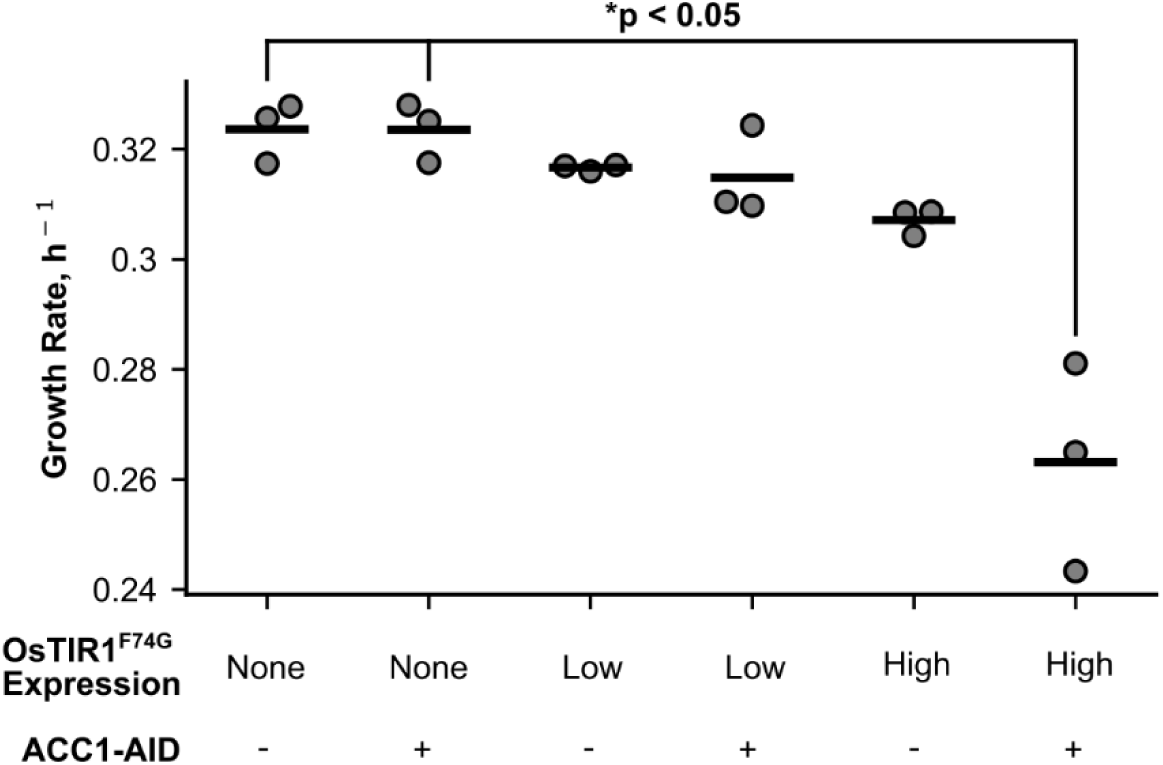
High basal degradation of Acc1-AID leads to growth defects. Growth rates of wild-type and strains expressing either OsTIR1^F74G^ alone or together with ACC1-mCherry-AID which is either expressed at a low (REV1 promoter) or high (ADH1 promoter) level. Growth rates from three biological replicates are shown for all strains. Fits for the determination of the individual growth rates per strain are shown in **Supplementary Figure 4**. Significance was determined using a Kruskal Wallis test followed by Dunn’s post-hoc test with Benjamini-Hochberg correction (p-value < 0.05, *).

In addition to the ACC1-AID strains, we constructed and analysed a strain in which TUP1, a non-essential gene encoding a transcriptional regulator, is AID-tagged and OsTIR1^F74G^ is expressed from a promoter of intermediate strength (i.e. the RNR2 promoter). In absence of auxin, we found that this strain strongly flocculated, forming a stable sediment at the bottom of the culture tube in liquid media. A control strain, only expressing TUP1-AID without OsTIR1^F74G^, did not exhibit any flocculation phenotype under the same growth conditions, demonstrating that the basal Tup1-AID degradation must be responsible for the flocculation phenotype. This observation is in line with earlier work, which reported flocculation to occur in TUP1 mutants^30–32^. The finding of the TUP1-AID flocculation phenotype due to basal degradation provides another reason for care when AID-tagging proteins.

## Discussion

In this work, we investigated the effect of OsTIR1^F74G^ expression levels on basal degradation of AID-tagged proteins as well as on the auxin-induced degradation dynamics. We found that basal degradation of AID-tagged GFP is dependent on OsTIR1^F74G^ expression. The degree of basal degradation ranges from a ∼4% using a low-strength promoter (pREV1) to up to ∼35% for high-strength promoters (pALD1, pRPL18B, pADH1). Furthermore, we showed that the rate of GFP-AID degradation upon 5-Ph-IAA addition is also dependent on OsTIR1^F74G^ expression, with a more than 5-fold difference between the minimal and maximal degradation rates. We further demonstrated that the auxin analogues NAA and 5-Ph-IAA can both be used to degrade GFP-AID in combination with OsTIR1^F74G^, although NAA-induced degradation exhibits slower dynamics than 5-Ph-IAA-induced degradation, and with NAA the steady state GFP-AID concentrations are only low when OsTIR1^F74G^ is highly expressed. Finally, we showcased that basal degradation of yeast proteins can lead to phenotypic changes. Basal degradation of AID-tagged Acc1 in a strain with high OsTIR1^F74G^ expression levels slowed down growth. Moreover, basal degradation of AID-tagged Tup1 in a strain with intermediate OsTIR1^F74G^ expression induced a flocculation phenotype consistent with a partial loss of Tup1 function.

As our work shows, even the optimized OsTIR1^F74G^ version does not prevent basal degradation of the AID-tagged protein in budding yeast. We suspect that this degradation might have been left unnoticed in the previous report where less quantitative spotting assays were used^2^. The degradation is consistent with the fact that the interaction of OsTIR1 with its cognate AID-sequence is only stabilised by auxin-binding, but not induced^14,15^. Along these lines, increasing OsTIR1^F74G^ levels simply increases the probability of an OsTIR1^F74G^-AID interaction, leading to protein degradation, even in the absence of auxin. Furthermore, the issue of basal degradation in the AID system might also be a feature of budding yeast, since budding yeast naturally produces IAA^33,34^.

In the present study, we assessed the degradation dynamics of cytosolic, highly expressed GFP-AID not fulfilling a function in yeast. Both the degradation dynamics and the steady-state level of a AID-tagged protein likely can be different for other proteins. Factors that will influence these values include the protein production rates, stability and general accessibility (e.g. cytosolic or within an organelle). For instance, genes residing in the mitochondria were reported to be less responsive to AID-induced degradation than cytosolic proteins^12^. Furthermore, the production of many proteins is regulated through processes on the genomic, transcriptional, and translational level^27^, also involving feedback mechanisms^27,35^, which could lead to counteraction of the induced protein degradation. Therefore, the choice of suitable OsTIR1^F74G^ expression levels will likely need to be adjusted to specifics of target proteins.

The use of the auxin degron system requires a careful balance between basal degradation and the required rate of protein degradation upon auxin addition. In cases where rapid degradation is required, high OsTIR1^F74G^ expression is necessary for fast degradation dynamics. However, when targeting essential proteins, or proteins whose concentrations should remain largely unaltered in the absence of auxin, elevated OsTIR1^F74G^ expression may lead to undesired basal degradation of the AID-tagged protein. If through variation of expression strength of OsTIR1^F74G^ no suitable balance between basal degradation and induced degradation dynamics can be achieved, more advanced dynamic control over OsTIR1^F74G^ levels and activity is needed, e.g. by the use of inducible OsTIR1^F74G^ expression^17^ or of specific OsTIR1 inhibitors^16^. Yet, such measures will make the perturbation system more complex, as both the induction of OsTIR1^F74G^ expression and the inhibition of OsTIR1^F74G^ are dynamic processes themselves.

Overall, the auxin degron system has proven to be a powerful research tool for conditional and temporal perturbation of biological systems^2,12^. Here, by systematically characterising important aspects such as basal degradation and degradation dynamics of the AID system in budding yeast, we improved the practical usability of the system.

## Methods

### Strains and Cultivation

The prototrophic YSBN6 strain derived from S288C ^36^ was used as genetic background for all yeast strains created in this work (see **Supplementary Table 1** for an overview of the yeast strains). For cloning purposes, strains were grown at 30 °C on yeast extract peptone dextrose (YPD) agar plates and in liquid YPD media, with and without selective antibiotics, respectively. For the actual experiments, strains were grown in minimal yeast nitrogen base (YNB) medium without amino acids and 2% D-glucose, buffered at pH 5.5 using a 100 mM K-Phthalate-KOH buffer (referred to as YNB-medium below). To obtain exponentially growing cell cultures, single colonies grown on selective YPD agar plates were inoculated into 10 mL of liquid YNB medium in 100 mL conical flasks and grown overnight, at 30 °C and shaking at 300 rpm. The overnight culture was then diluted to a cell count of 1.0-2.5 ‧ 10^6^ mL^−1^ and grown to a cell count of 8.0-15.0 ‧ 10^6^ mL^−1^ during the day. Assuming a growth rate of 0.3 h^−1^, the over-day culture was subsequently diluted in YNB-medium to reach a cell count of 10.0-15.0 ‧ 10^6^ mL^−1^ the next day. The next day, these exponentially growing cultures were again diluted to a desired cell count and maintained at an exponential growth phase for the rest of the day by keeping their cell count below 20.0 ‧ 10^6^ mL^−1^ by dilution.

### Growth rate measurement and analysis

To measure the specific growth rate of strains, we prepared exponentially growing yeast cultures as described above and measured cell counts in regular time-intervals using a BD Accuri C6 flow-cytometer. We then linearly regressed the log-transformed cell density values using the sklearn.linear_model package^37^ implemented in python and derived the growth rate from the slope of the regression line. Statistical tests were conducted using the scipy. stats^38^ module and scikit.posthocs^39^ packages implemented in Python.

### Plasmid construction

For the construction of plasmids encoding the genetic elements required for the measurement of basal degradation, competent *E. coli* DH5α cells^40^ were used for transformation using a standard heat-shock method^41^. For an overview of plasmids and primers used and created in this study, refer to **Supplementary Table 2** and **Supplementary Table 3**, respectively. For plasmid design, we applied a modular cloning approach according to the Molecular Cloning Yeast Toolkit (MoClo-YTK) as described^23^. The MoClo-YTK makes use of basic part-plasmids that can be combined to higher order transcriptional units and multigene-plasmids via Golden Gate assembly for expression in *S. cerevisiae*. All Golden Gate assemblies were performed as described^23,42^. The correctness of all constructs was confirmed by sequencing.

Part-plasmids were constructed by a BsmBI Golden Gate assembly with part pYTK001 as an entry vector. For the construction of type-3 (coding sequence) part pYTK235, GFP-AID was amplified from pG2J^9^ using primer pair JF54-JF72 carrying type-3 part 5’-overhangs for subsequent Golden Gate assembly into pYTK001. Type 2 promoter part pYTK234 was constructed by amplification of P_TETO7__part1 and P_TETO7__part2 using primer pairs JF64-JF65 and JF66-JF67 from plasmid pB_teto7_sfGFP, respectively, to eliminate undesired type IIs restriction sites. The two PCR products P_TETO7__part1 and P_TETO7__part2 were subsequently inserted into pYTK001. For construction of type-4 (terminator) part pYTK236, the CYC1 terminator was PCR-amplified using primer pair JF68-JF69 carrying part 4-type 5’-overhangs for subsequent Golden Gate assembly into pYTK001. Inserts for Type-2 (promoter) part pYTK238 (ADH1 promoter) and type 3 parts pYTK240 (OsTIR1^F74G^ coding sequence), pYTK258 (non-codon-optimized OsTIR1^F74G^ coding sequence) and pYTK237 (sfGFP coding sequence) were synthesized and cloned into the canonical pYTK001 entry vector by BsmBI Golden Gate assembly. For codon optimization of OsTIR1^F74G^, we used the GeneSmart^TM^ codon optimization tool offered by Genescript^43^. Part pYTK241 was constructed by amplification of the mCherry coding sequence from pG23A with primer pair JF73-JF74 and subsequent cloning into pYTK001 by BsmBI Golden Gate assembly. Part pYTK242 was constructed by amplification of S3-5GA-AID^71–114^ from pG2J with primer pair JF75-JF76 and subsequent cloning into pYTK001 by BsmBI Golden Gate assembly. Parts pYTK253 and pYTK254 were constructed by amplification of the 3’-HO and 5’-HO region from YSBN6 genomic DNA using primer pairs JF99-JF100 and JF97-JF98, respectively, and subsequent cloning into pYTK001 by BsmBI Golden Gate assembly.

Cassette plasmids were constructed by BsaI Golden Gate assembly into pYTK095 entry vector. The pJF01 cassette plasmid was constructed by BsaI Golden Gate assembly of parts pYTK002, pYTK013, pYTK235, pYTK236 and pYTK067 into the pYTK095 entry vector. Cassette plasmid pJF02 was constructed by BsaI Golden gate assembly of parts pYTK002, pYFK067, pYTK234, pYTK237 and pYTK236 into the pYTK095 entry vector. Cassette plasmid pJF05 was constructed by BsaI Golden gate assembly of parts pYTK067, pYTK061, pYTK013, pYTK002, pYTK042, pYTK241 and pYTK242 into the pYTK095 entry vector. The cassette plasmids, containing promoters of varying strengths and either codon-optimized or non-codon optimized OsTIR1^F74G^ versions, all contained the common parts pYTK003, pYTK054, pYTK072 together with pYTK240 and pYTK258, respectively. To increase or decrease the OsTIR^F74G^ codon-optimized expression strength, the common parts were assembled together with pYTK238, pYTK017, pYTK018, pYTK023, pYTK027 and pYTK022 to obtain cassette plasmids pJF04, pJF10, pJF11, pJF12, pJF13 and pJF14, respectively. To express non-codon-optimized OsTIR1^F74G^, pYTK238 was assembled together with the common parts and pYTK258 to obtain plasmid pJF30. The cassette plasmid pJF25 encoding the GFP-AID control was constructed by BsaI Golden Gate assembly of parts pYTK253, pYTK013, pYTK235, pYTK236 and pYTK254 into the pYTK095 vector.

Multigene plasmids were constructed by BsmBI Golden Gate assembly into pYTK101/pDN4 as previously^42^. Multigene plasmids containing GFP-AID and the codon optimized OsTIR1^F74G^ version expressed from different promoter strengths were assembled using the GFP-AID containing part pJF01 together with pJF04, pJF10, pJF11, pJF12, pJF13 and pJF14 to obtain pJF06, pJF15, pJF16, pJF17, pJF18 and pJF19, respectively. Multigene plasmids pJF07, pJF22 and pJF23 containing sfGFP and the codon optimized OsTIR1^F74G^ version expressed from ADH1, RNR2 or REV1 promoter were assembled using the sfGFP containing part pJF02 together with pJF04, pJF12 or pJF13, respectively. The multigene plasmid containing GFP-AID and the non-codon-optimized OsTIR1^F74G^ version expressed from the ADH1 promoter was assembled with pJF01 together with pJF30 to obtain pJF34.

For construction of the plasmid containing sgRNA targeting the HO-locus and ACC1-C-terminus in YSBN6, we followed a protocol based on the yeast molecular cloning kit as previously^23,42^. To construct the plasmid containing the sgRNA targeting the ACC1-C-terminus of YSBN6, we first annealed oligos JF03 and JF04 and cloned them into pYTK050 using a BsmBI Golden Gate assembly to create the ACC1-C-terminus-sgRNA part plasmid pYTK-RB1. Next, we performed a BsaI Golden Gate assembly with pYTK-RB1, pYTK003, pYTK068 and pYTK095 to create the ACC1-C-terminus-sgRNA cassette plasmid pYTK-RB2. Ultimately, we created the final Cas9_ACC1-C-terminus-sgRNA multi-gene plasmid pYTK-RB3 by BsmBI Golden Gate assembly using pYTK-DN1, pYTK-DN2, pYTK-RB2, and pYTK-DN5. To construct the plasmid containing the sgRNA targeting the TUP1-C-terminus of YSBN6, we first annealed oligos JF140 and JF141 and cloned them into pYTK050 using a BsmBI Golden Gate assembly to create the TUP1-C-terminus-sgRNA part plasmid pJF48. Next, we performed a BsaI Golden Gate assembly with pJF48, pYTK003, pYTK068 and pYTK095 to create the TUP1-C-terminus-sgRNA cassette plasmid pJF60. Ultimately, we created the final Cas9_TUP1-C-terminus-sgRNA multi-gene plasmid pJF76 by BsmBI Golden Gate assembly using pYTK-DN1, pYTK-DN2, pJF60, and pYTK-DN5.

### Strain construction

All strains used in this study were constructed using CRISPR-Cas9 genome editing as described^42^. All constructs were integrated into the HO-locus of YSBN6 and transformations were performed according to an established protocol^44^. We designed and cloned a single guide RNA (sgRNA) targeting the HO-locus or the ACC1 c-terminus using the e-crisp or CRISPOR web applications as described^42,45,46^.

To construct strains yJF01, yJF02, yJF10-yJF14, yJF20, yJF23 and yJF27, we linearized plasmids pJF06, pJF07, pJF15, pJF17-pJF19, pJF16, pJF23, pJF34 and pJF22, respectively, by PCR with primers JF81-JF82. The linear fragment for construction of yJF15 was obtained by NotI digestion of pJF25 and subsequent purification of the digestion product. The purified PCR and digestion products were then integrated into the HO-locus of a YSBN6 wild-type strain by CRISPR-Cas9 genome editing using plasmid pYTK-AK3 which contains the Cas9 expression cassette and HO-sgRNA cassette. To construct strain yJF07, we linearized plasmid pJF05 with primer pair JF77-JF104 and used the purified PCR product to tag the c-terminal ACC1-sequence of yJF02 strain by CRISPR-Cas9 genome editing with plasmid pYTK-RB3 which contains the Cas9 expression cassette and Acc1-c-terminal-sgRNA cassette.

To construct strain yJF08, we linearized plasmid pJF05 with primer pair JF77-JF104 by PCR and used the purified PCR product to tag the c-terminal ACC1-sequence of a YSBN6 wild-type strain by CRISPR-Cas9 genome editing with plasmid pYTK-RB3 which contains the Cas9 expression cassette and Acc1-c-terminal-sgRNA cassette. Strain yJF21 was constructed by linearization of pJF23 using primers JF81-JF82 and subsequent integration into the HO-locus of yJF08 using plasmid pYTK-AK3 which contains the Cas9 expression cassette and HO-sgRNA cassette.

To construct strain yJF43 and yJF44, we linearized plasmid pJF05 with primer pair JF155-JF167 by PCR and used the purified PCR product to tag the c-terminal TUP1-sequence of a YSBN6 wild-type strain and yJF27, respectively, by CRISPR-Cas9 genome editing with plasmid pJF76 which contains the Cas9 expression cassette and TUP1-c-terminal-sgRNA cassette.

The correctness of constructed strains was confirmed by PCR and DNA sequencing of the DNA region of interest.

### Flow cytometry measurements and data analysis

For cell count and cellular GFP-fluorescence measurements, we used a BD Accuri C6 flow-cytometer. We measured the GFP-AID fluorescence in the FL1-A channel using a 488-nm laser for excitation and 533 nm ± 30 nm to detect emission. Following the GFP-fluorescence measurement, we exported the raw, single cell measurements as FCS-files using the BD CFlow Plus analysis software and analysed the data using the FlowCal package^47^ implemented in Python. After transforming the FL1A-data in the fcs-file to arbitrary fluorescence units (a.u.), we filtered out unspecific events likely stemming from cell debris or non-cellular particles by applying a density gate based on the FSC-H and SSC-H channels, capturing 90% of the total event counts. To account for differences in individual cell size and compute a GFP-AIP concentration estimate per cell, we divided each single cell’s FL1-A value by its respective FSC-A value, which is a proxy for cell size^48^. We then computed the median of these single-cell GFP-AID concentration values which approximates to the GFP-AID concentration (*GFPC*) value per strain. We background corrected the *FL*-values of each sampled strain GFPC_*sample*_ and the *GFPC*_*control*_-values of a control strain that constitutively expresses GFP-AID, but does not express OsTIR1^F74G^ (see strain yJF15 in **Supplementary Table 2**) by subtracting the GFP-AID (auto)-fluorescence *GFPC*_*wwt*_ of the YSBN6 wild type strain from both *GFPC*_*sample*_ and *GFPC*_*control*_. To enable comparison of *GFPC*-values across experiments, we then calculated GFPC_*norm*_by normalizing the background corrected *GFPC*_*sample*_ values to the background corrected *GFPC*_*control*_ values:

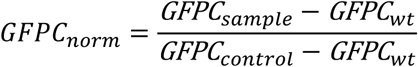

### Measurement of basal degradation

To determine basal degradation, we measured the fluorescence of 1.0-2.0·10^4^ single cells taken from exponentially growing yeast cultures using a BD Accuri C6 flow cytometer at a flow rate of 35 μL·min^−1^ and a core size (diameter of sample flow in between the sheath laminar flow) of 16 µm. The *GFPC*_*norm*_ values per technical replicate were determined as described above and used to compute the mean *GFPC*_*norm*_ across the technical replicates which ultimately represents the *GFPC*_*norm*_ value per biological replicate.

### Measurement and analysis of auxin-induced degradation dynamics using method 1

To determine the auxin-induced degradation dynamics using method 1, exponentially growing cells were diluted to a cell count of 2.0 × 10^6^ mL^−1^ and allowed to grow for 1h at 30 °C and 300 rpm before adding 5-Ph-IAA or 1-NAA to a final concentration of 5 µM or 500 µM, respectively. For each strain, we took fluorescence measurements just before and in time intervals after addition of auxin using a BD Accuri C6 flow-cytometer at a flow rate of 35 μL·min^−1^ and a core size of 16 µm. At each sampling timepoint, we took 1.0-2.0·10^4^ single cell fluorescence measurements. Since the measurement frequency and timespan varied between different replicate experiments, we pooled the time course measurements of different experimental replicates per strain together. We normalised individual measurements as described above and thereby obtained the normalised, single cell fluorescence *GFPC*_*norm*_-values at each timepoint for each strain. To estimate degradation rates and GFP-AID concentration at steady state, we fit a one-phase exponential decay function to the normalised time course median fluorescence measurements for each strain. We applied non-linear least squares regression using the trust region reflective method^49^ implemented in scipy^38^ with the following model function:

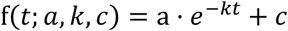

where t is the time after addition of auxin in *min*, *a* is the unitless initial cellular GFP-AID fluorescence intensity before addition of auxin, *k* is the degradation rate constant in min^−1^and c is the unitless cellular GFP-AID fluorescence intensity when a steady state between GFP-AID degradation and production has been reached. Parameter bounds were chosen as follows:

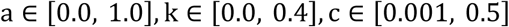

The lower parameter bounds constrain the fit to non-negative values for *a* and *k*. We chose c > 0.001 since a steady state is always higher than 0 fluorescence units due to the continuous expression of GFP-AID at steady state. The upper parameter bounds were constrained for *c* ≤ 0.5 which was a value higher than all observed steady state GFP-AID concentration based on visual inspection of the time course data, for *k* ≤ 0.04 since the observed values of *k* were well below 0.4 in each strain, and for *a* ≤ 1.0 since the initial GFP-AID measurements should not be higher than the control strain without any apparent basal degradation.

To estimate uncertainty of parameter estimates per strain, we employed a bootstrapping approach. From each set of fluorescence time course measurements per strain, we drew 1000 bootstrap samples with replacement using the numpy.random module^50^. Next, we fitted the one-phase exponential decay function to each of the resampled bootstrap samples per strain as described above. To mitigate the effect of randomness inherent to bootstrap sampling, we repeated the bootstrapping procedure five times without setting a random seed and aggregated the resulting parameter estimates.

### Measurement and analysis of auxin-induced degradation dynamics using method 2

To determine the auxin-induced degradation rate using method 2, exponentially growing cells were diluted to a cell count of 0.5-1.0 · 10^6^ mL^−1^ and a final volume of 1.8 mL in YNB-medium in 96-deep-well plates and continuously measured with a flow cytometer using a flow rate of 10 μL · min^−1^ and a core size of 8 µm. After 5 minutes of continuous measurement, the flow was stopped for less than a minute and 5-Ph-IAA was added to a final concentration of 5 µM and mixed before the flow was started again for another 30-40 minutes. For testing the dependence of different auxin concentrations on the degradation rate of GFP-AID in the strain expressing pREV1, we added auxin to a final concentration of 2.5 µM, 5.0 µM, 10 µM, 20 µM and 50 µM. Following flow cytometry acquisition, the data was filtered and normalised as described above to obtain different time course measurements with a high time-resolution across the sampling timepoints. A one-phase exponential decay function was then fitted to the measurements as described above to derive the degradation rate *k* from the resulting fits.

## Acknowledgements

Funding from the Dutch Research Council (NWO) to M.H. (grant number VI.C.192.003) is acknowledged.

## Supplementary Tables

**Supplementary Table 1:**
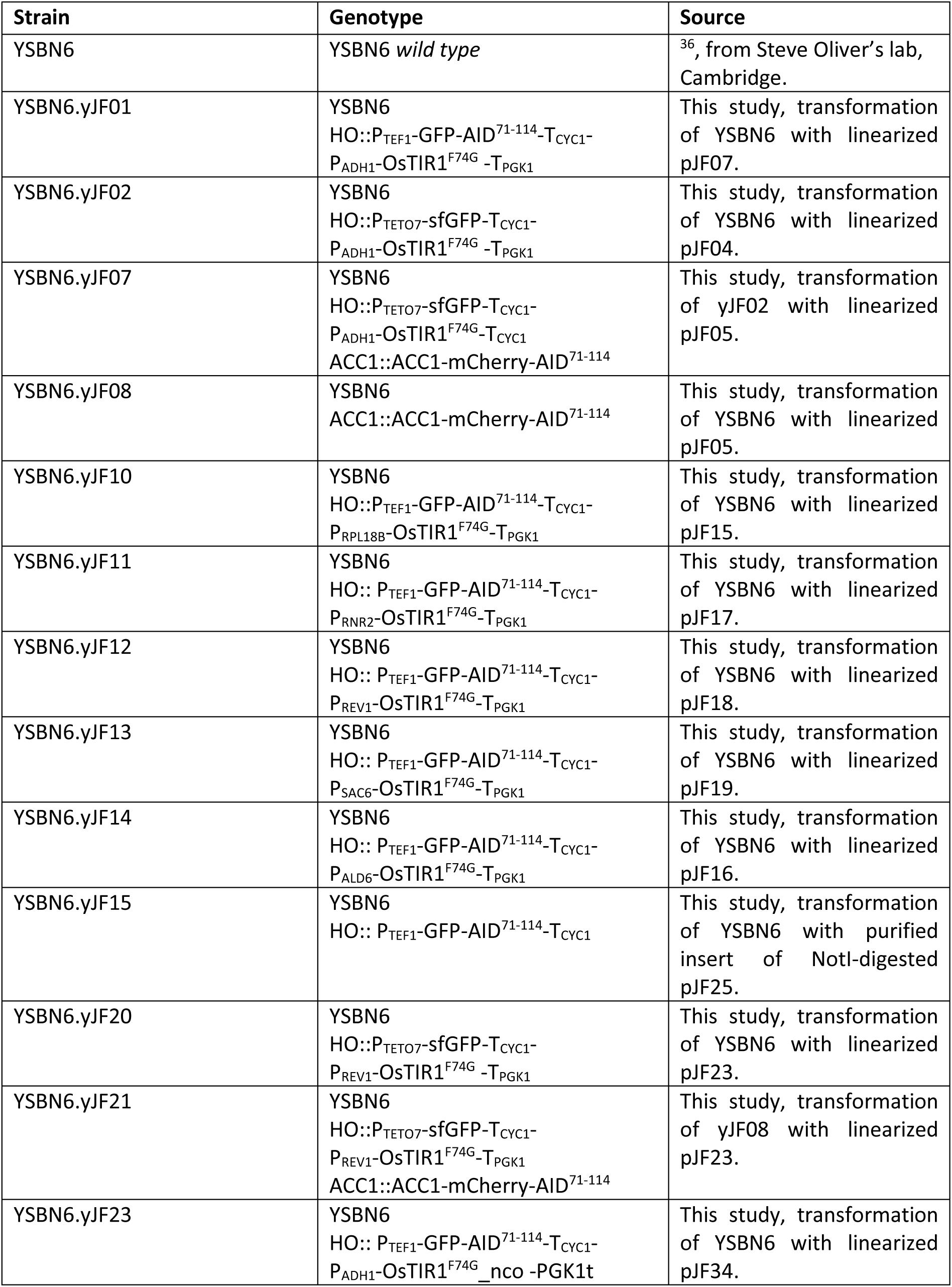
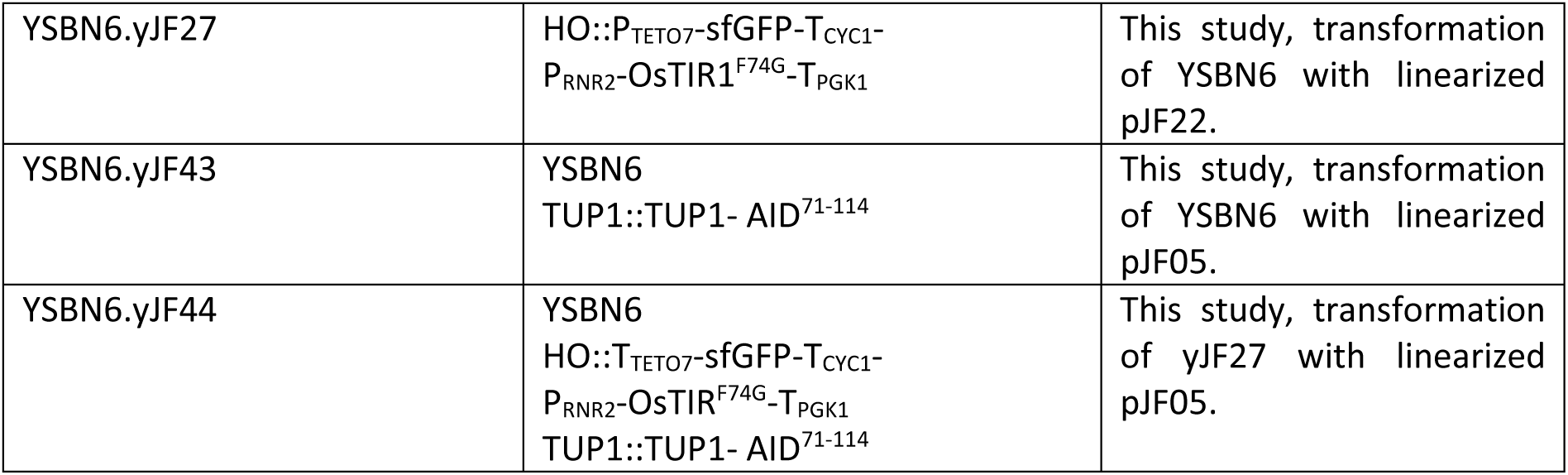
Yeast strains created in this study. Nco: non-codon optimized.

**Supplementary Table 2:**
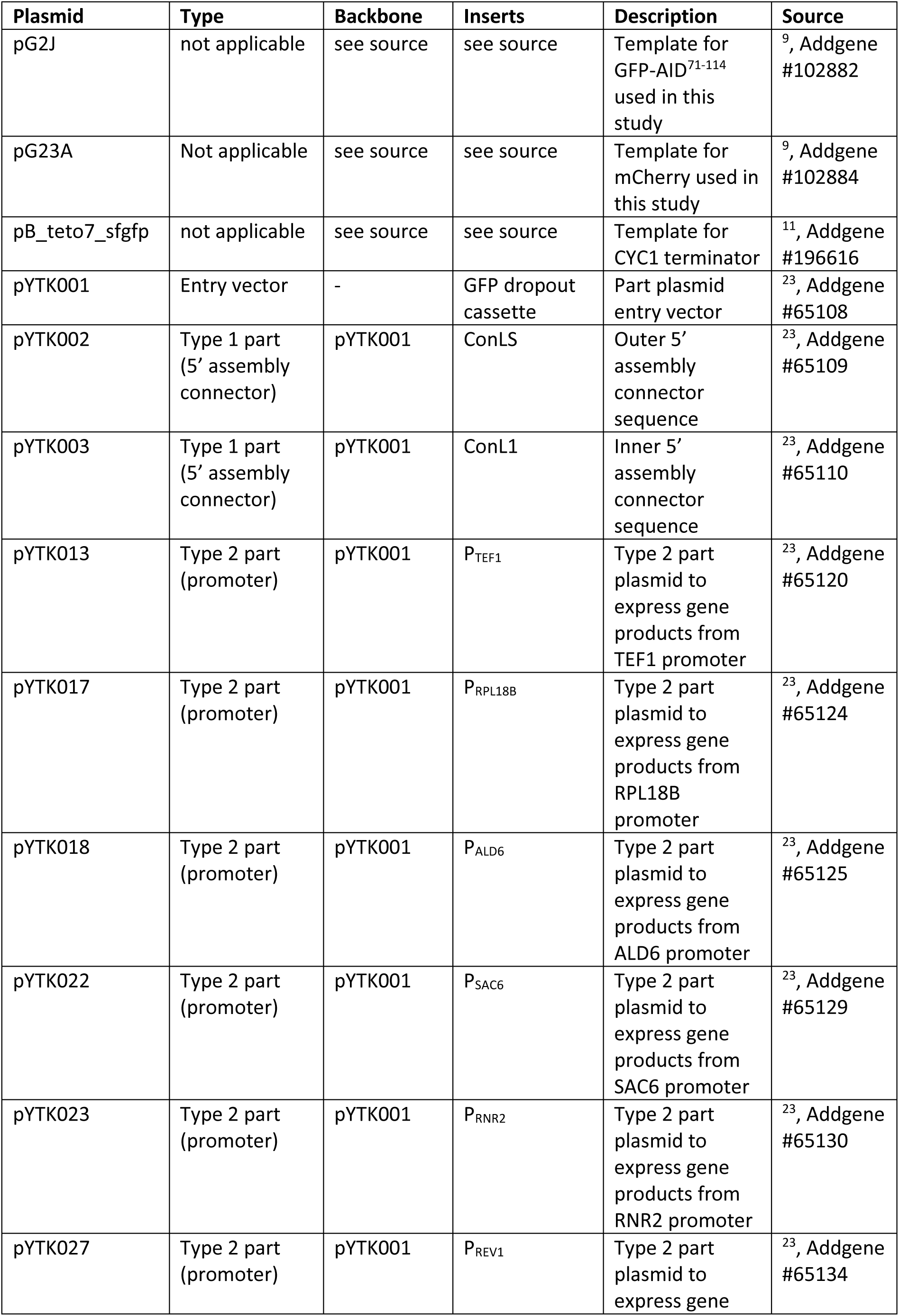
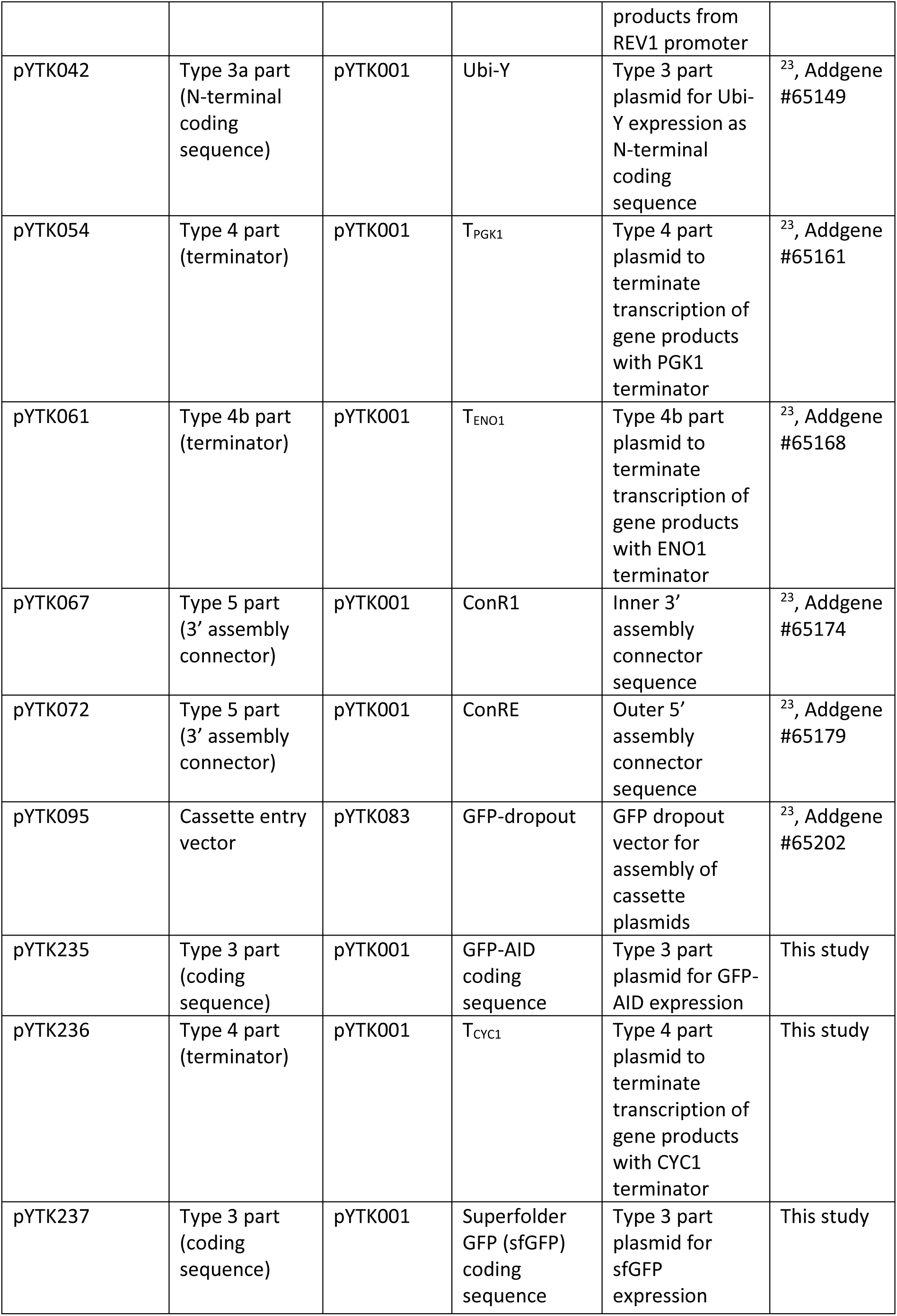
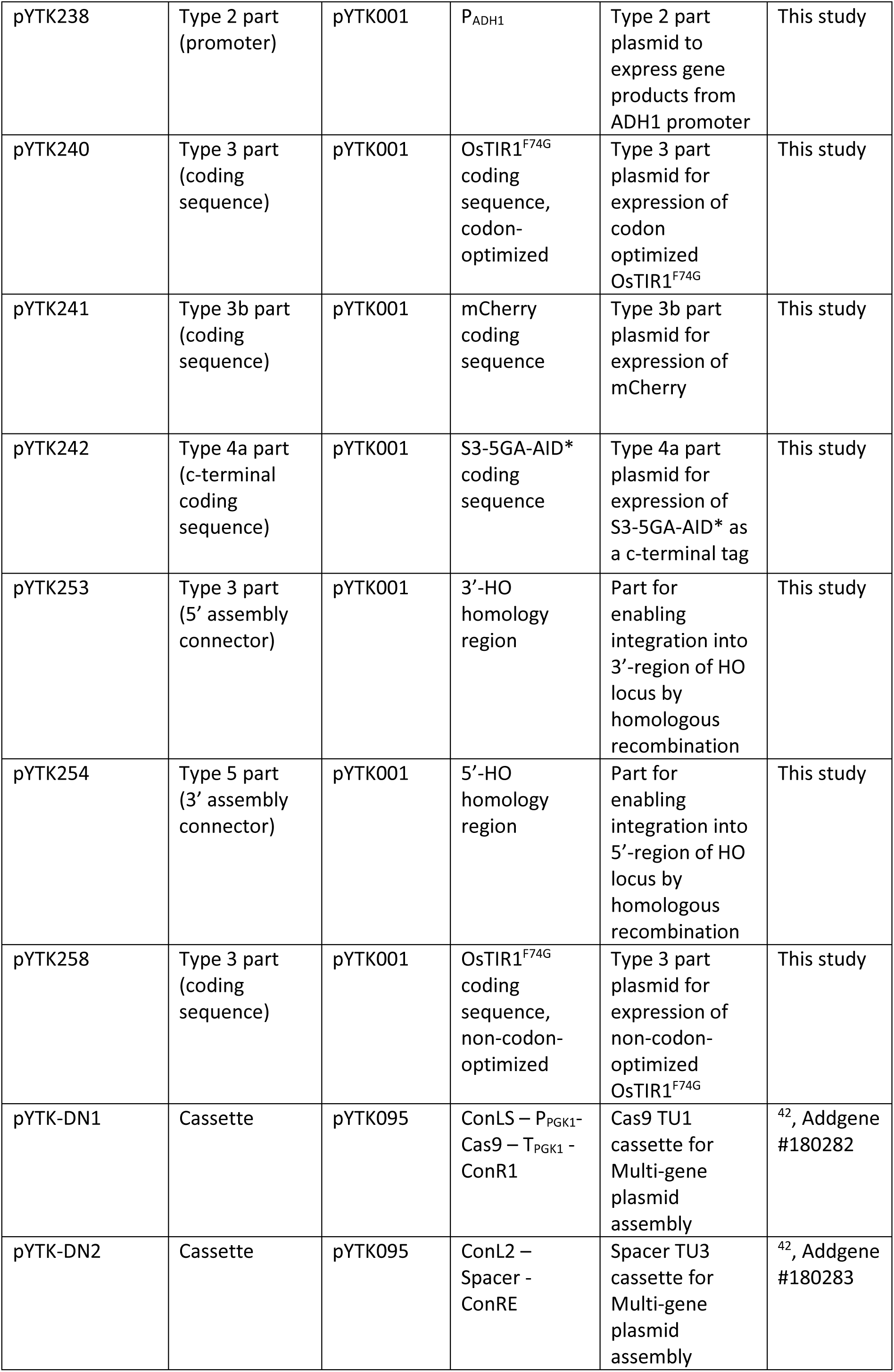
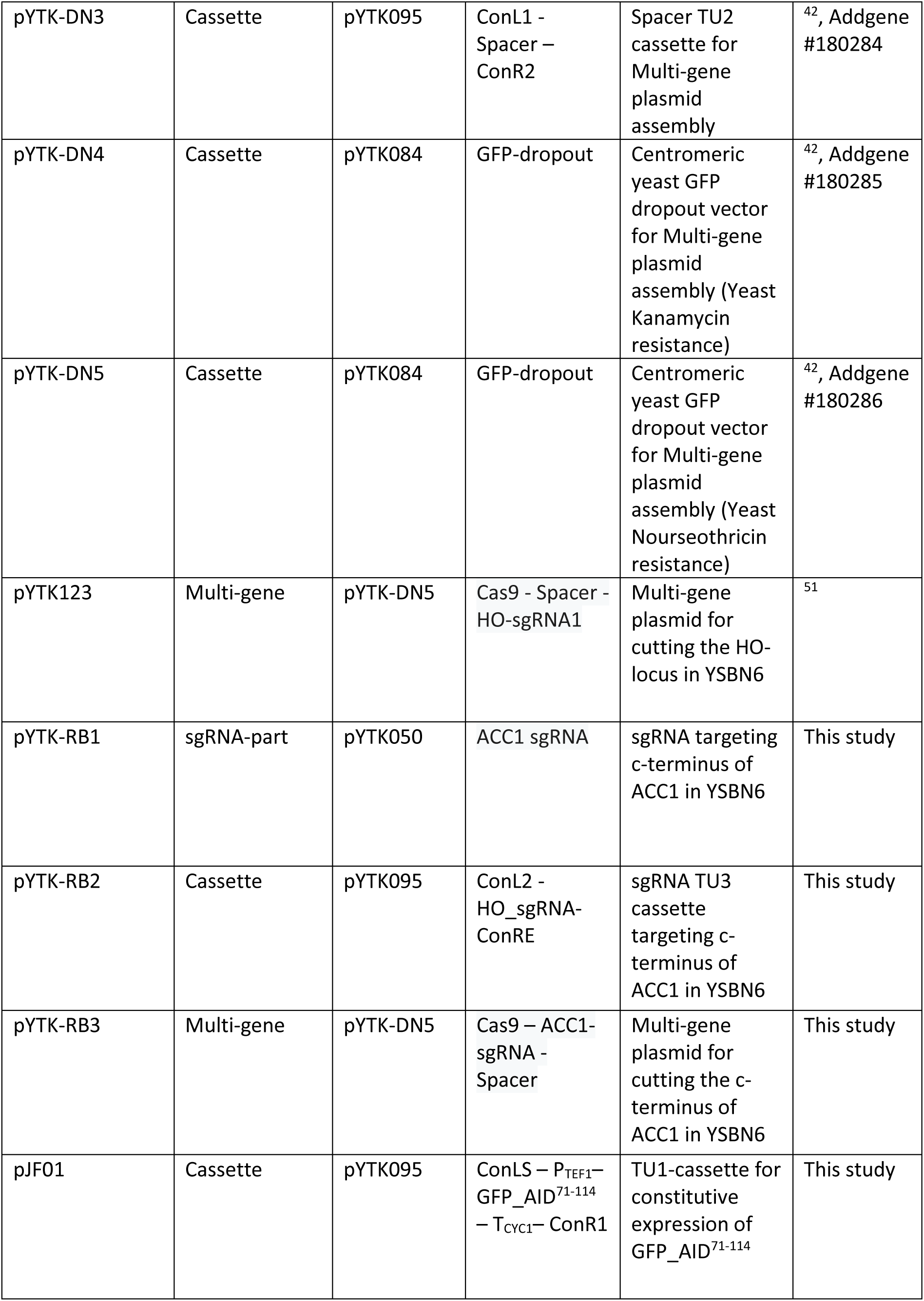
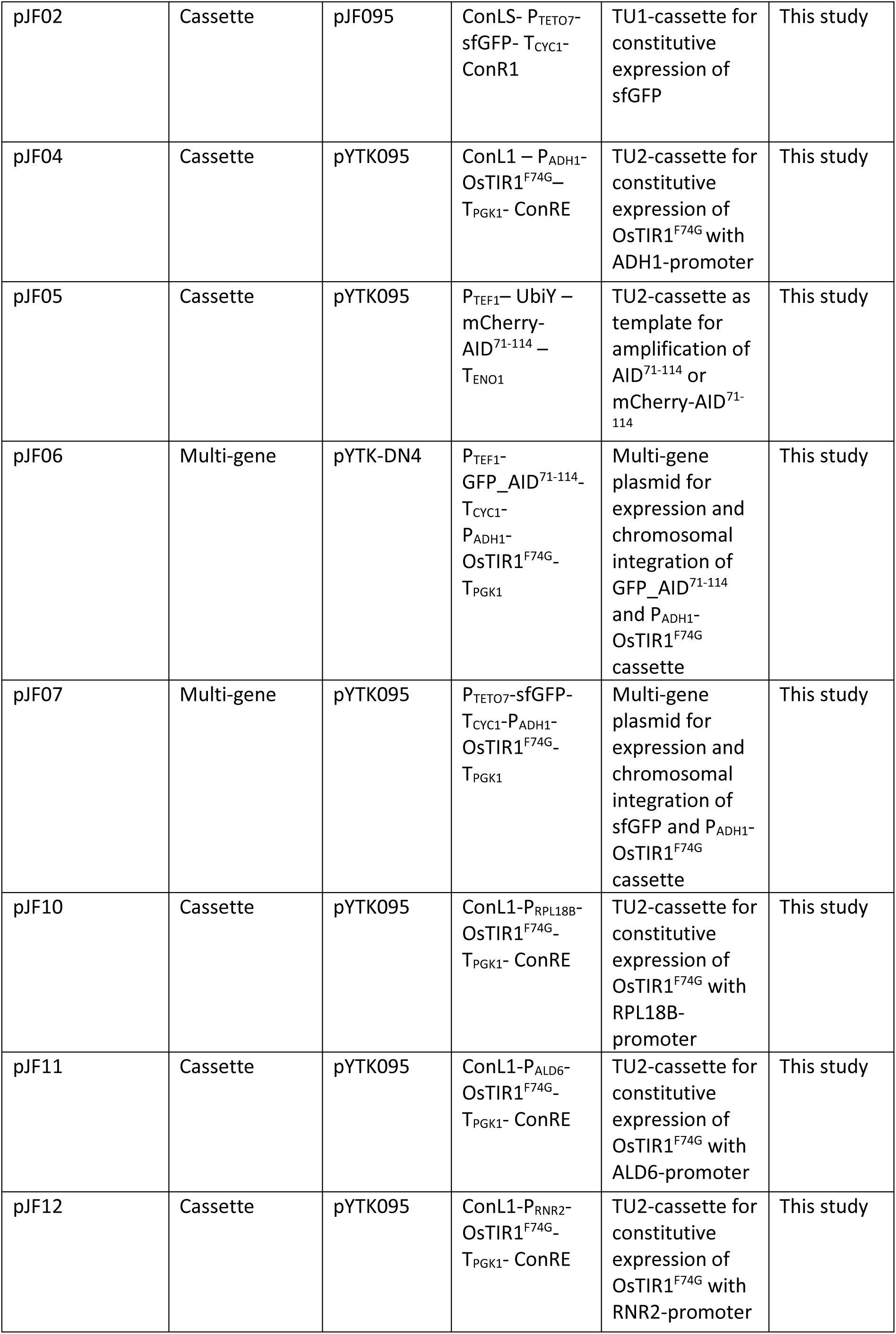
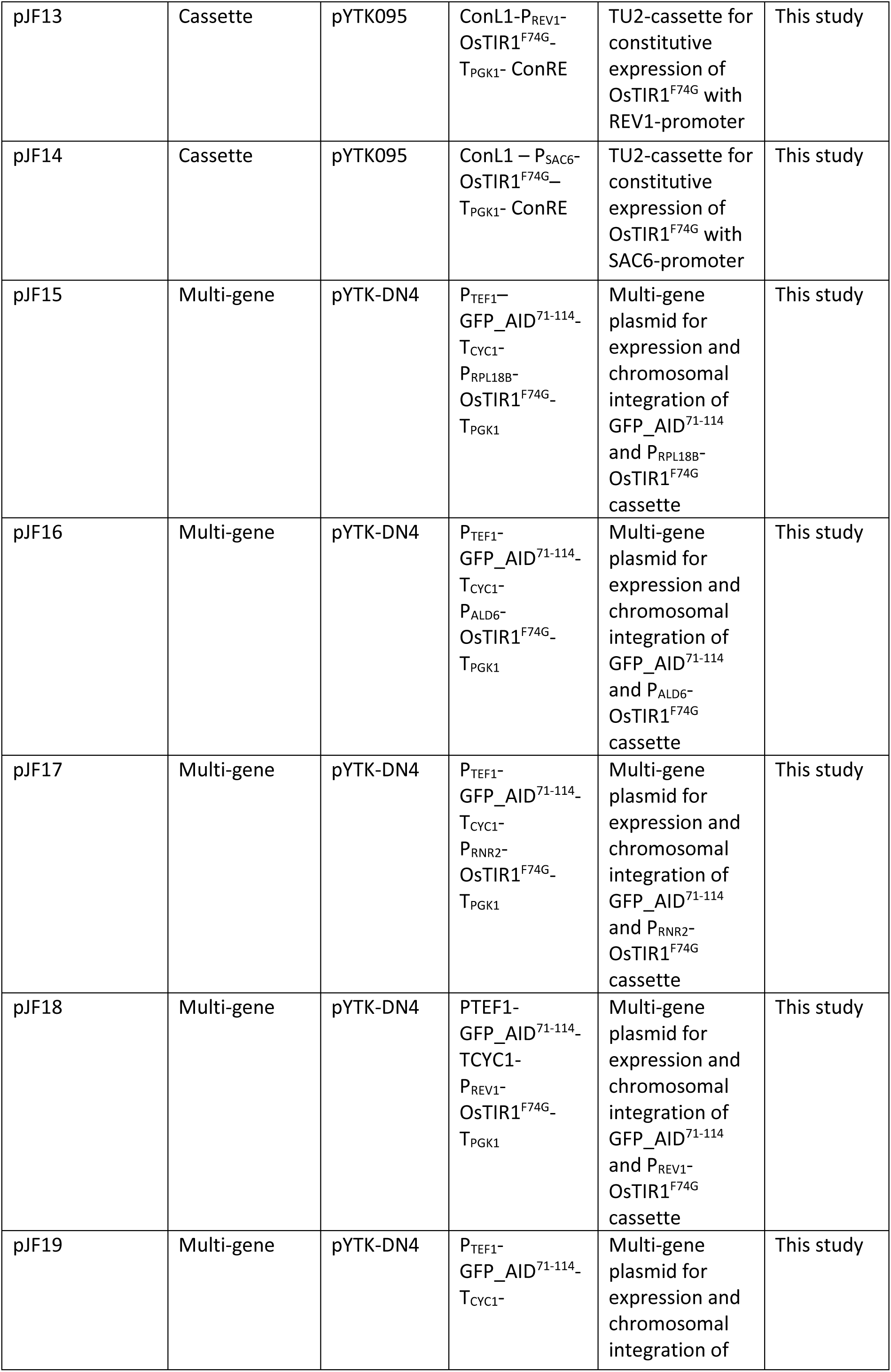
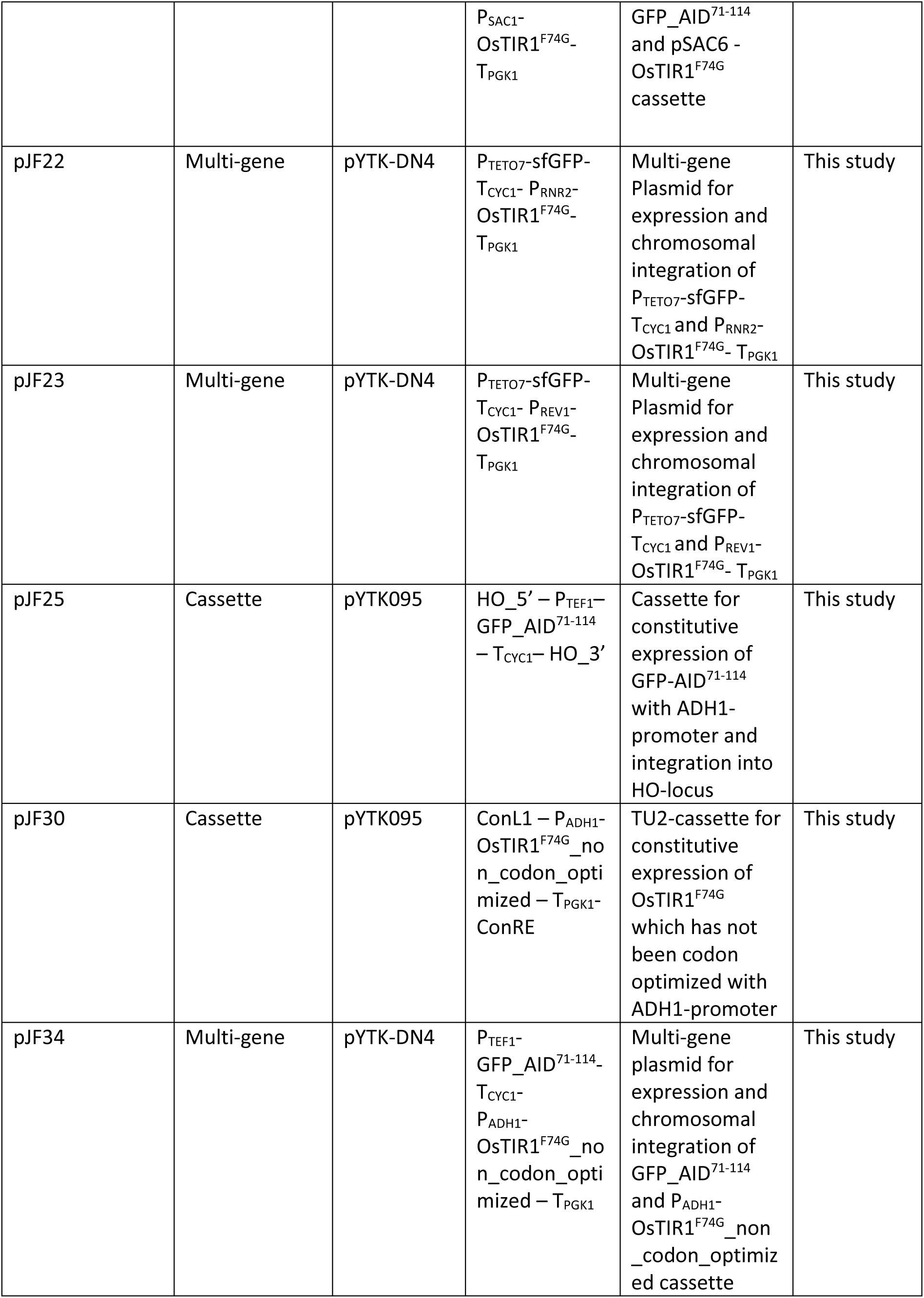
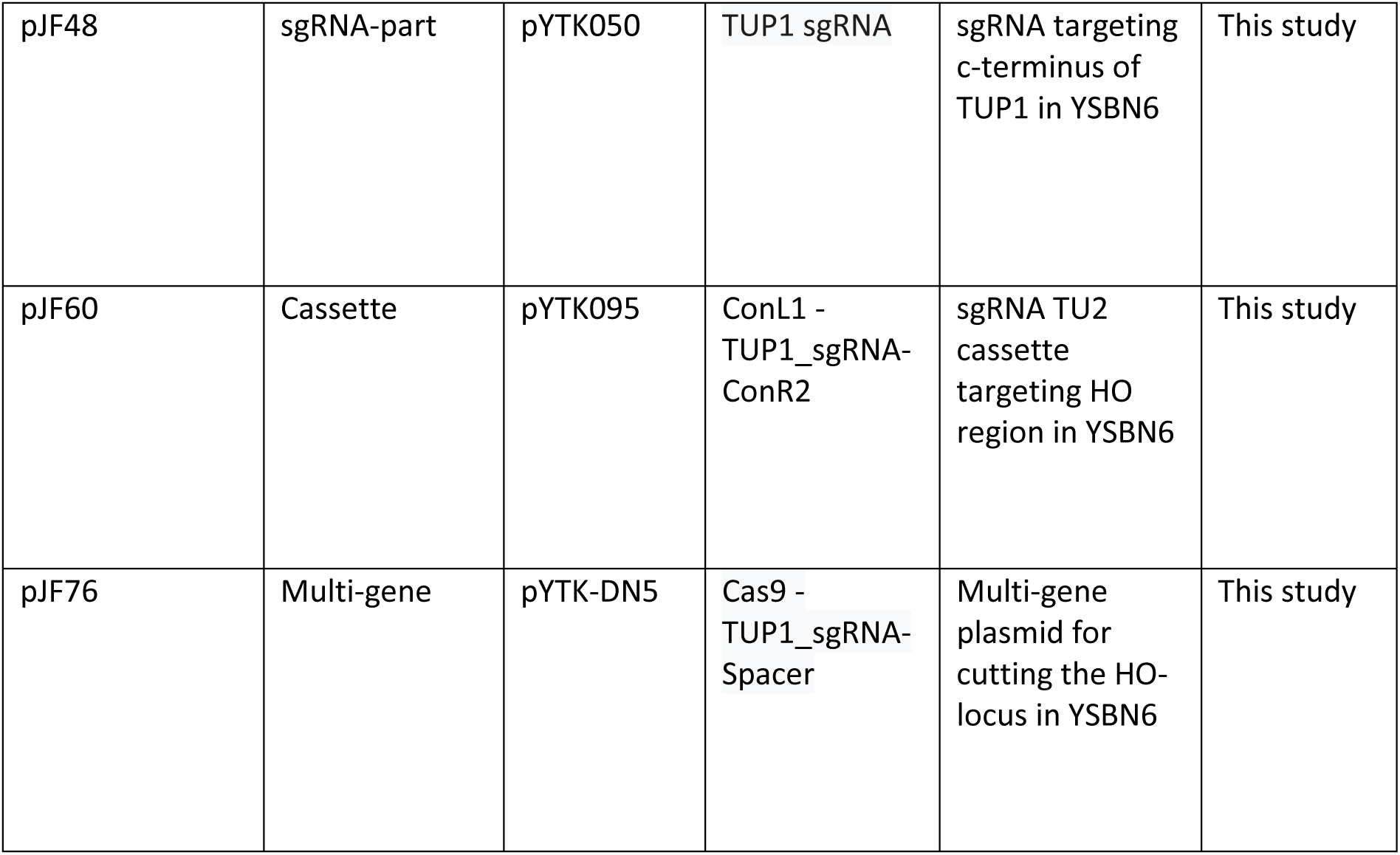
Plasmids used for the construction of all recombinant yeast strains.

**Supplementary table 3:**
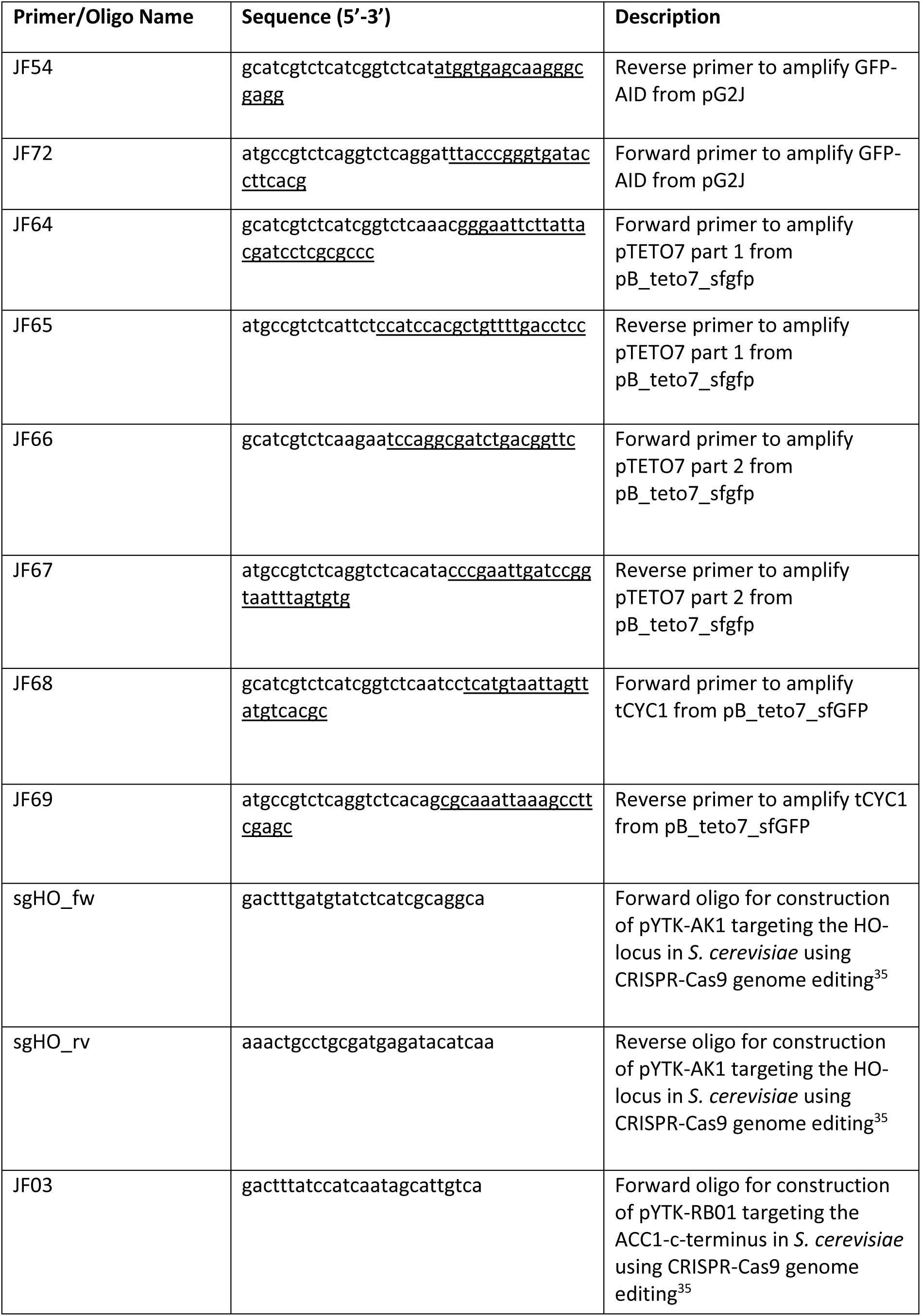
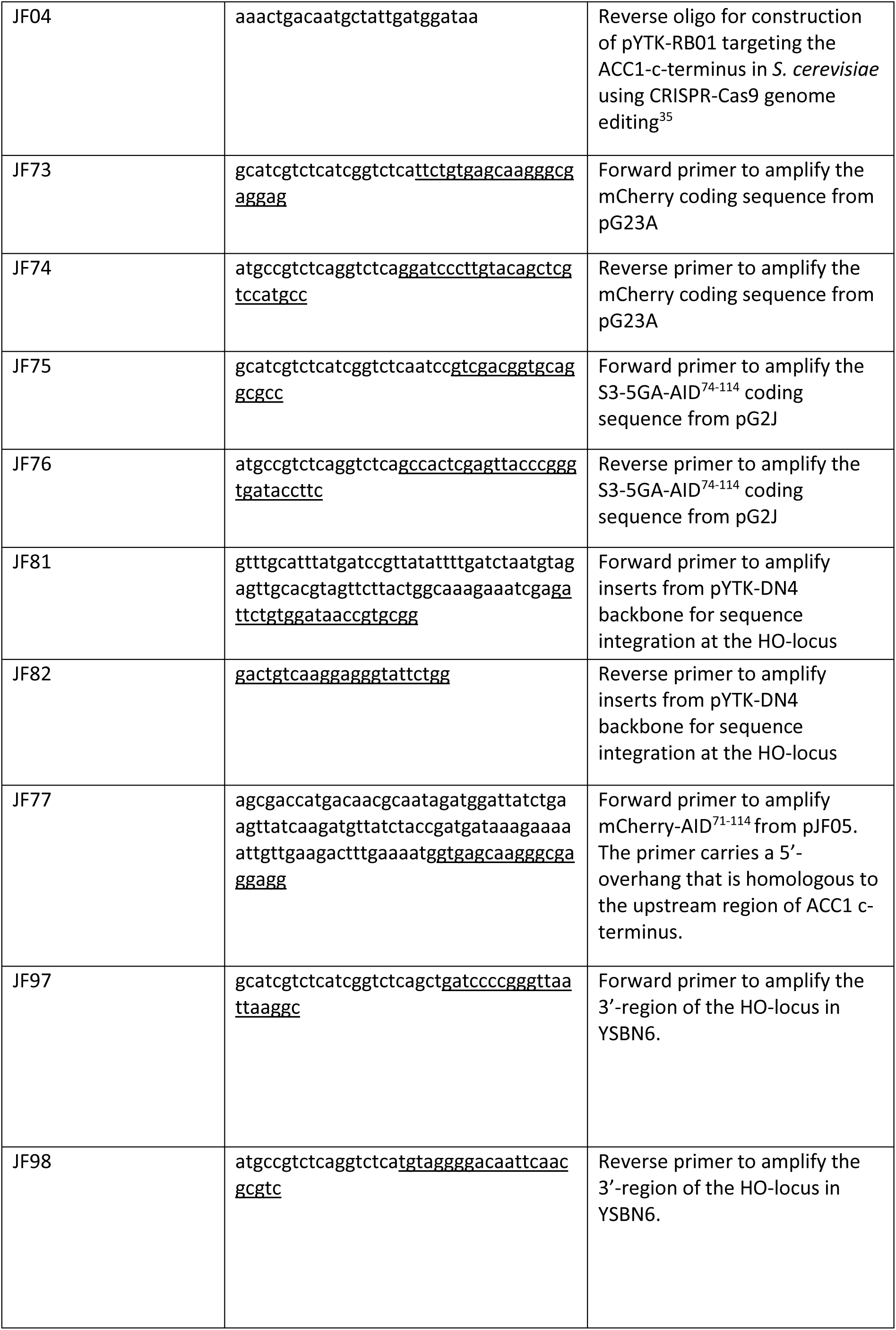
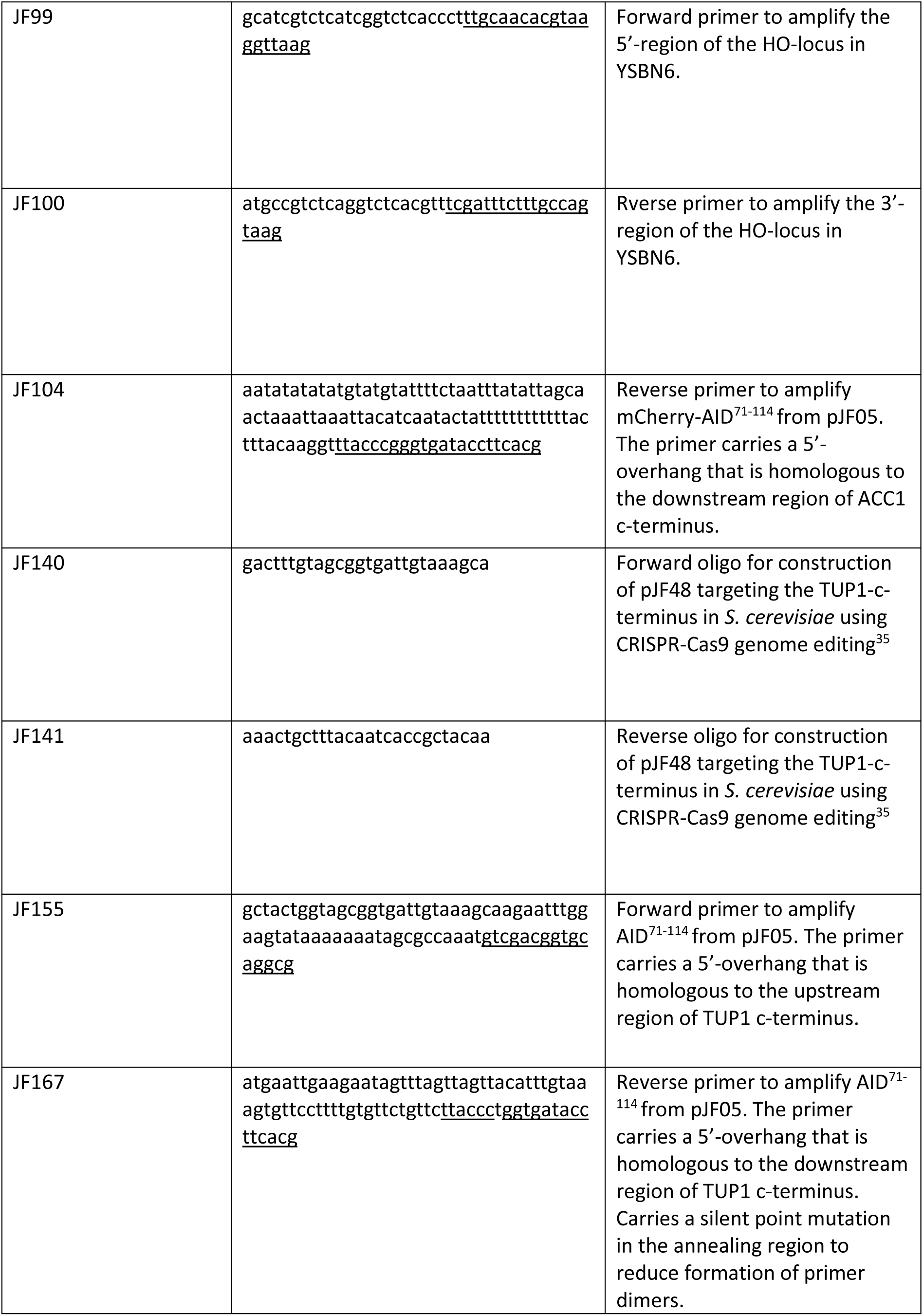
Primers and oligos used in this study. Underlined sequences denote the DNA-binding regions of primers.

**Supplementary table 4:**
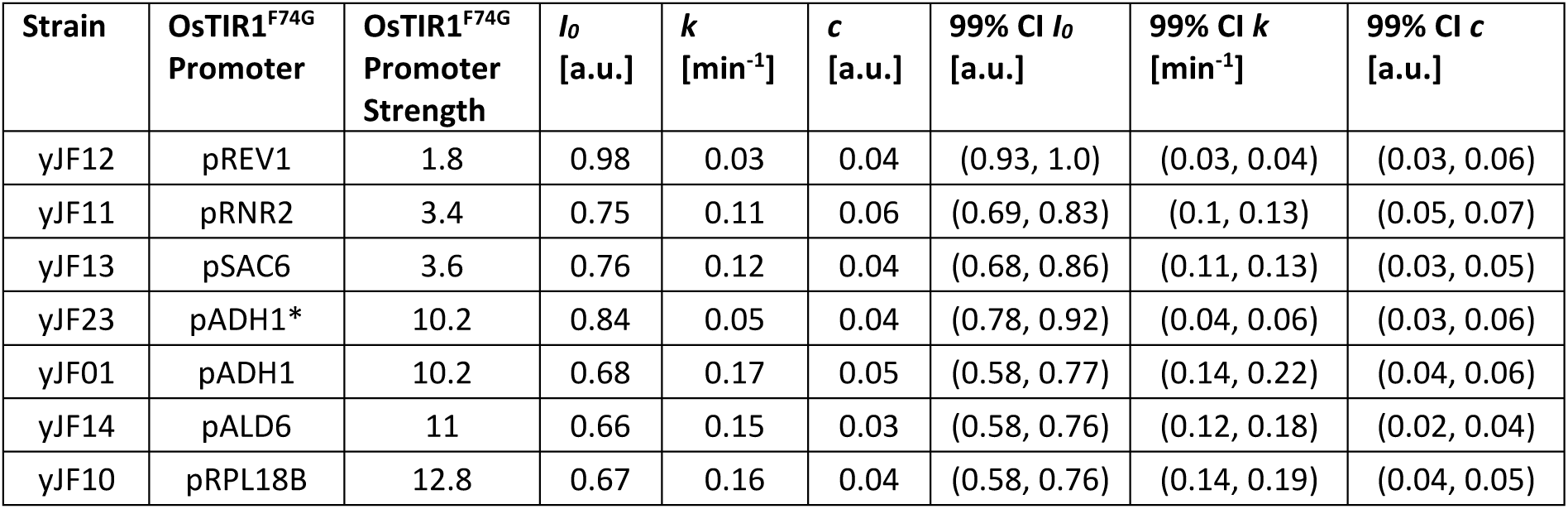
Parameter estimates and their uncertainties for GFP-AID degradation dynamics induced with 5-Ph-IAA using method 1.

**Supplementary table 5:**
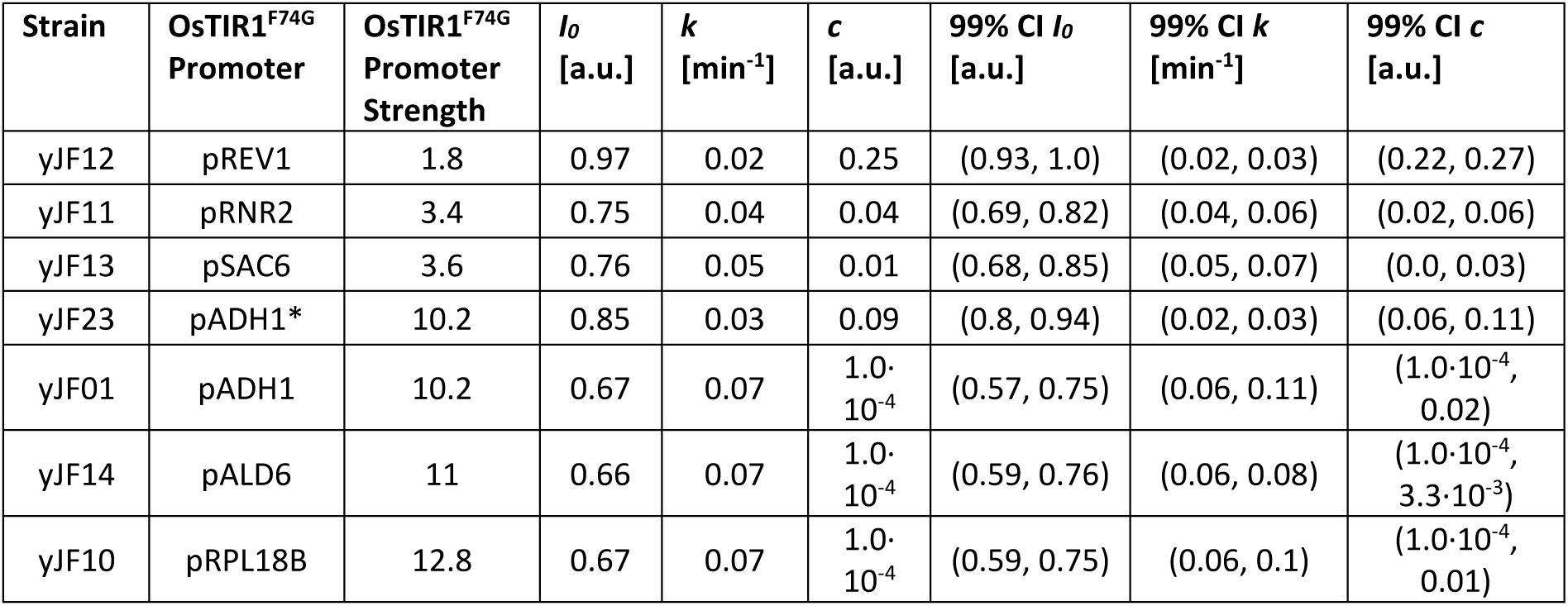
Parameter estimates and their uncertainties for GFP-AID degradation dynamics induced with NAA using method 1.

---

**Supplementary Figure 1:**
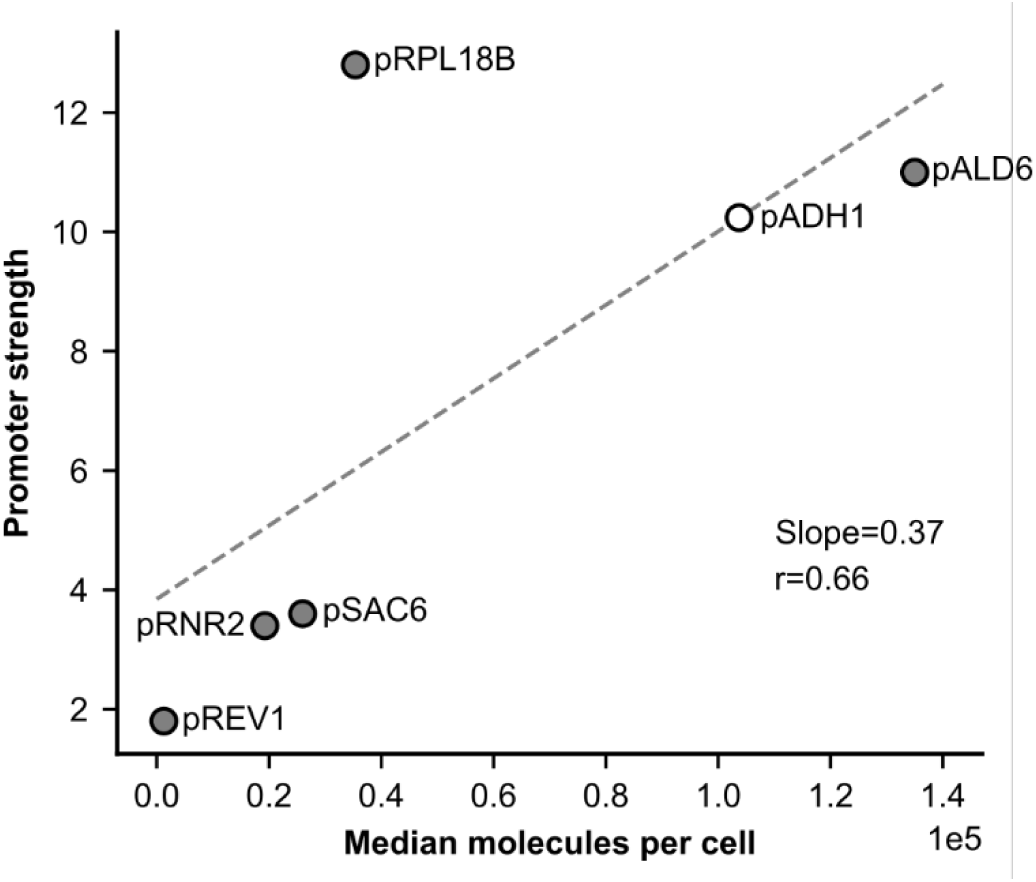
Determination of OsTIR1^F74G^ promoter strengths. Promoter strengths from Lee *et al.*, 2015^23^ based on the mRuby2 fluorescence measurements presented in Figure 3 of the same publication plotted against the median single molecule number per cell of each protein belonging to the respective promoter^25^. In this study, we used the promoter strengths as indicator of OsTIR1^F74G^ expression strengths. Since the promoter strength of the ADH1 promoter was not present in the publication of Lee *et al.*^23^, we estimated the ADH1 promoter strength by linear regression based on the promoter strengths and single molecule protein numbers of Rev1p, Rnr2p, Sac6p, Ald6p and Rpl18bp^25^ assuming that the reported, relative protein levels roughly correlate with the expression strength of each promoter. Based on this regression, together with the determined promoter strengths according to Lee *et al.*, 2015, we placed the ADH1 promoter strength between the promoter strength of pALD6 and pRPL18B. Closed symbols represent promoters whose expression strengths were reported in Lee *et al.*^23^, whereas the open symbol represents the estimated promoter strength of pADH1. The dashed line is the regression line of linear regression of the promoter strengths and protein molecules per cell (slope=0.37, r=0.66).

**Supplementary Figure 2:**
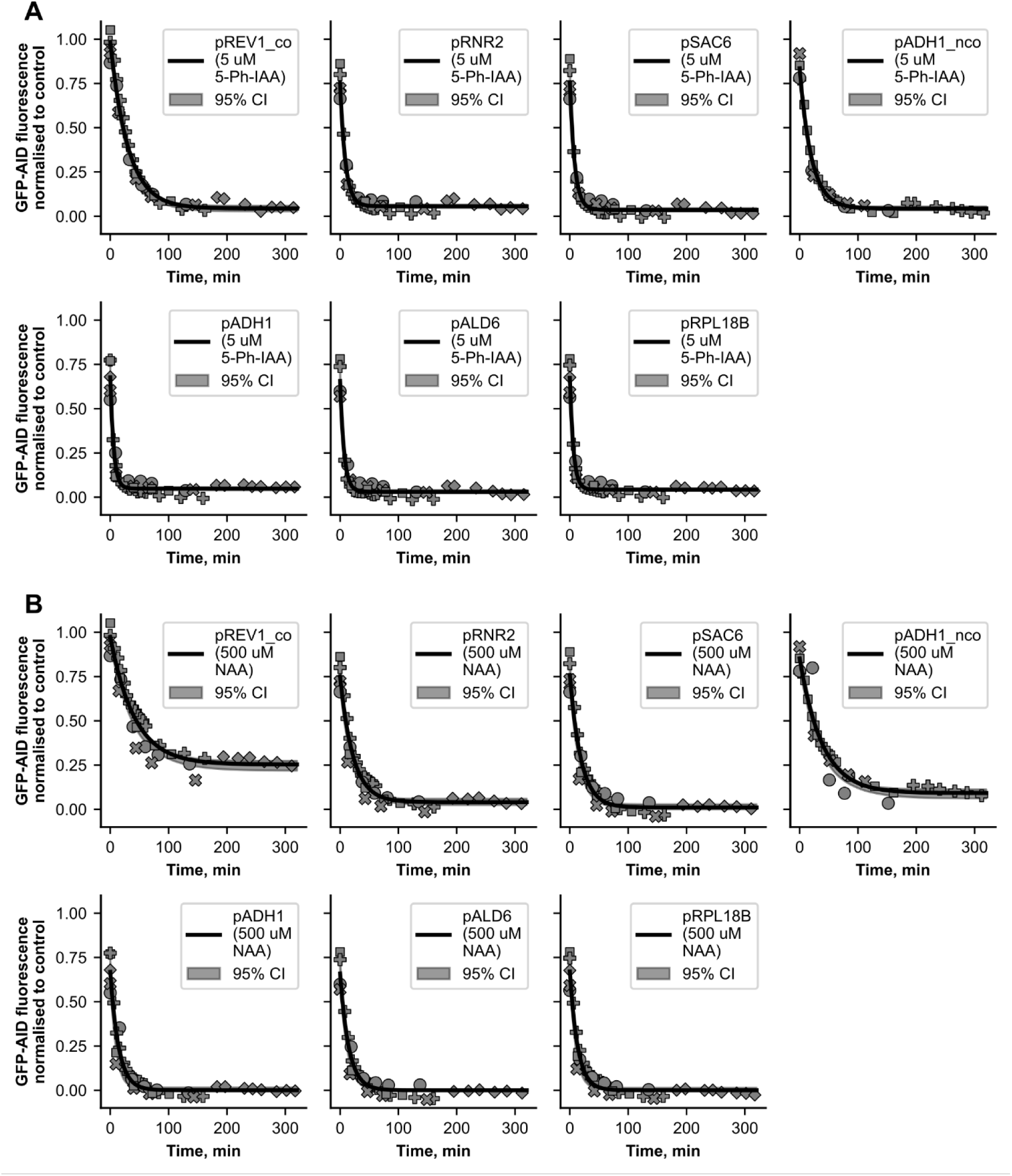
Exponential decay fits for each strain after addition of 5-Ph-IAA (A) and NAA (B). The different symbols indicate measurements belonging to different biological replicates. The shaded area around the model fit indicate the 95% confidence interval of the fit obtained by bootstrapping.

**Supplementary Figure 3:**
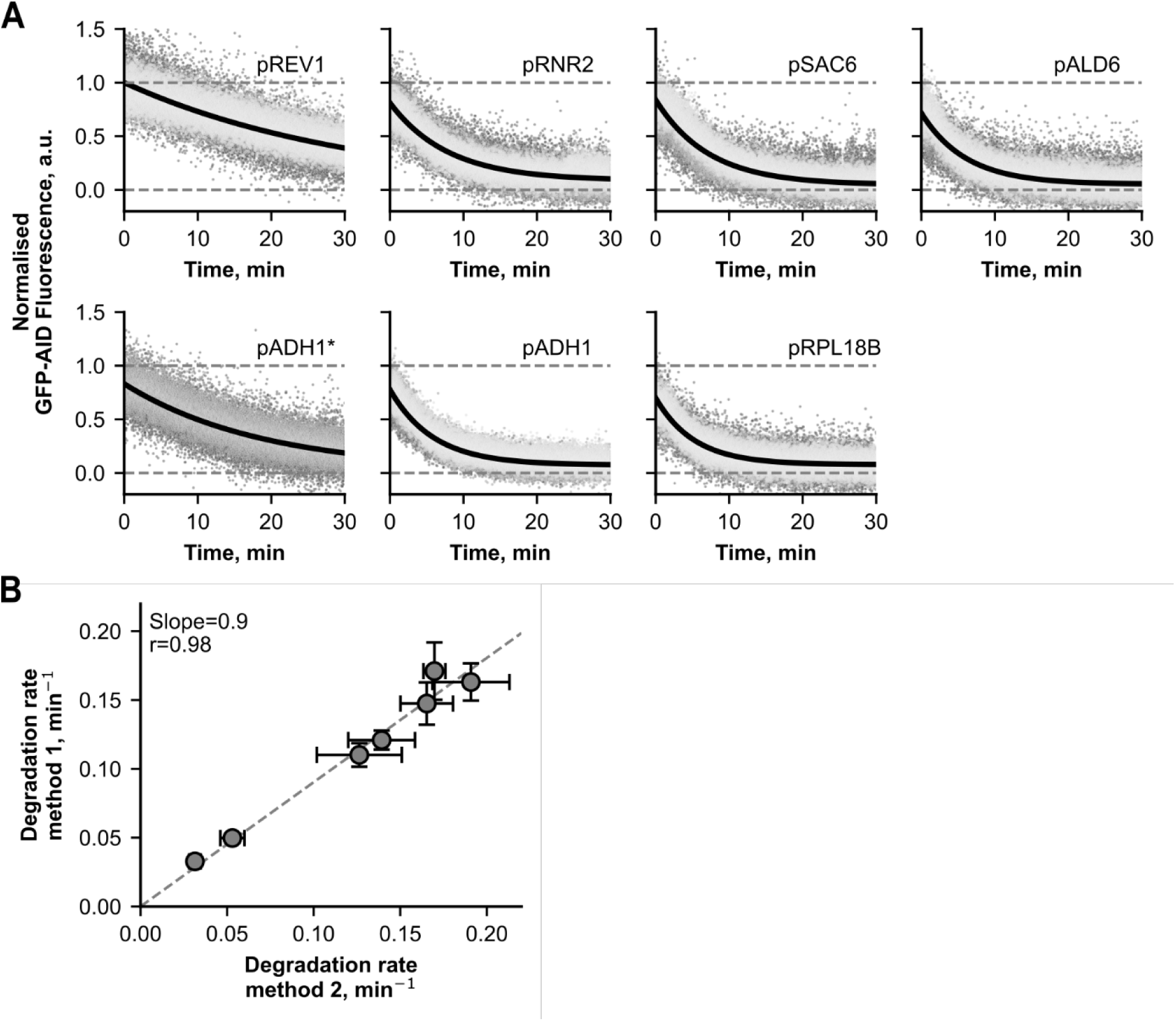
Degradation rates determined with an alternative method match the observed degradation rates per strain. **(A)** Fits of the exponential decay function to data obtained from the alternative method 2 for determination of degradation rate k per strain. Different shades of gray indicate an individual replicate of the three biological replicates. **(B)** Comparison of the degradation rates k obtained by method 1 and the alternative method 2. Filled symbols denote strains expressing a codon-optimized OsTIR1^F74G^ gene and the empty symbol denotes the strain expressing non-codon optimized OsTIR1^F74G^. Error bars indicate standard deviations of the estimates of k for different replicates. The dashed line represents a linear regression of the degradation rates determined by method 2 against method 1 (slope=0.9, r=0.98).

**Supplementary Figure 4:**
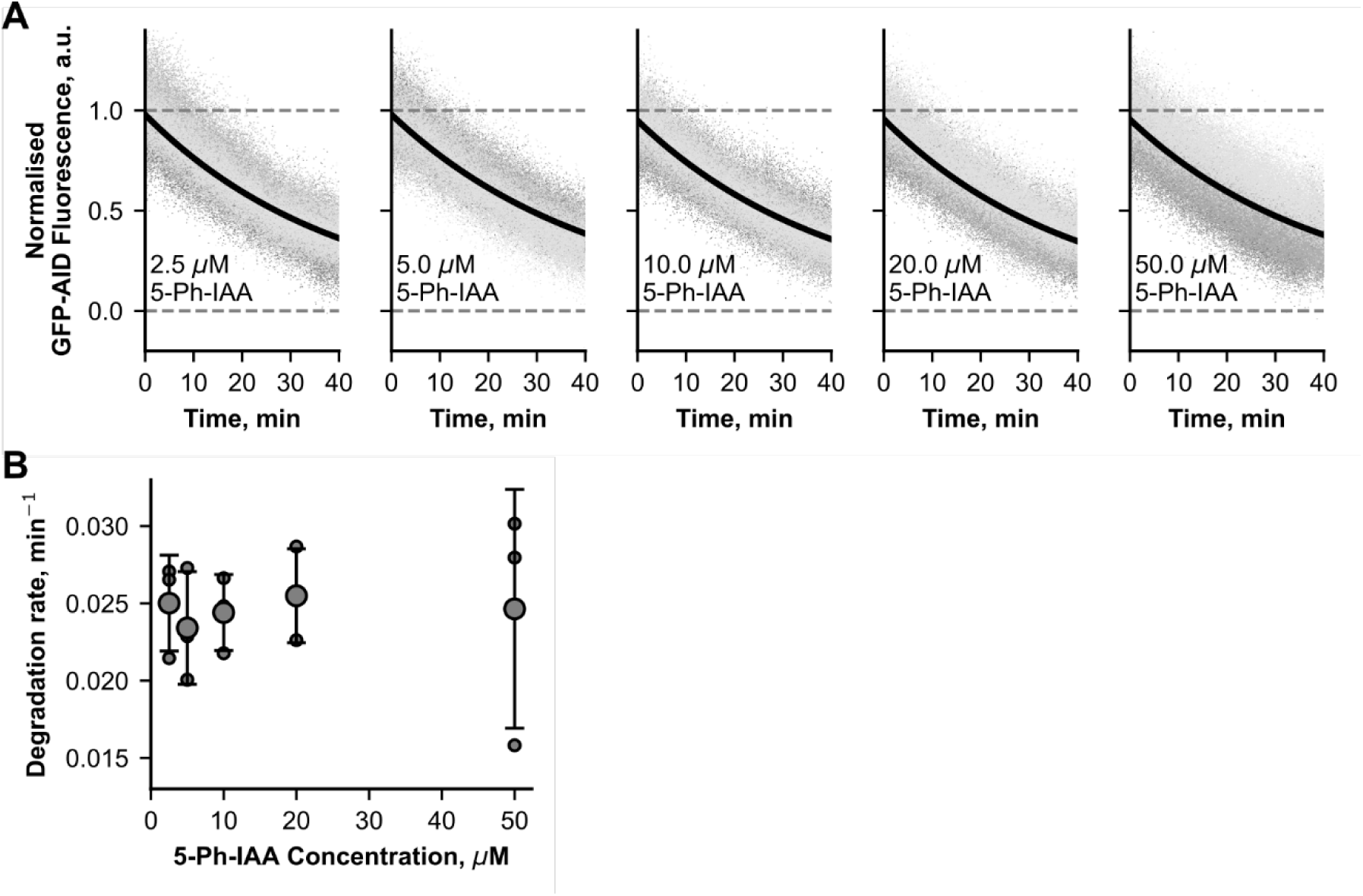
An increase in 5-Ph-IAA concentration does not increase the degradation rate of strain yJF12. **A** Individual fits of exponential decay functions to GFP-AID fluorescence signal after adding 5-Ph-IAA at different concentrations to a strain expressing OsTIR1^F74G^ from the lowest-strength promoter tested. In each plot, an exponential decay function is fitted to data from three different biological replicates. **B** Degradation rates of the individual fits shown in **A**. Small datapoints denote the degradation rates determined per replicate, while big datapoints represent the mean degradation rate of the three different biological replicates. Error bars denote the standard deviation across the replicates.

**Supplementary Figure 5:**
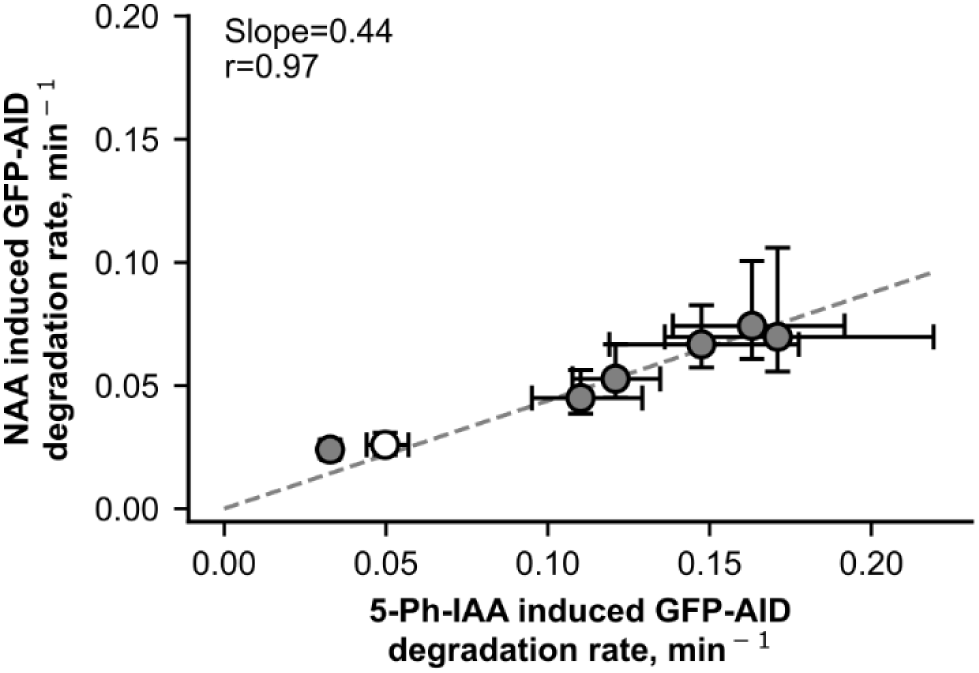
Comparison of NAA and 5-Ph-IAA induced degradation rates using method 1. Filled symbols represent strains with codon-optimized OsTIR1^F74G^, the empty symbol represents a strain expressing non-codon optimized OsTIR1^F74G^. Error bars indicate the 95% confidence interval for the estimates of the degradation rates. The dashed line describes a linear regression of NAA-induced degradation against 5-Ph-IAA induced degradation (slope=0.44 r=0.97).

**Supplementary Figure 6:**
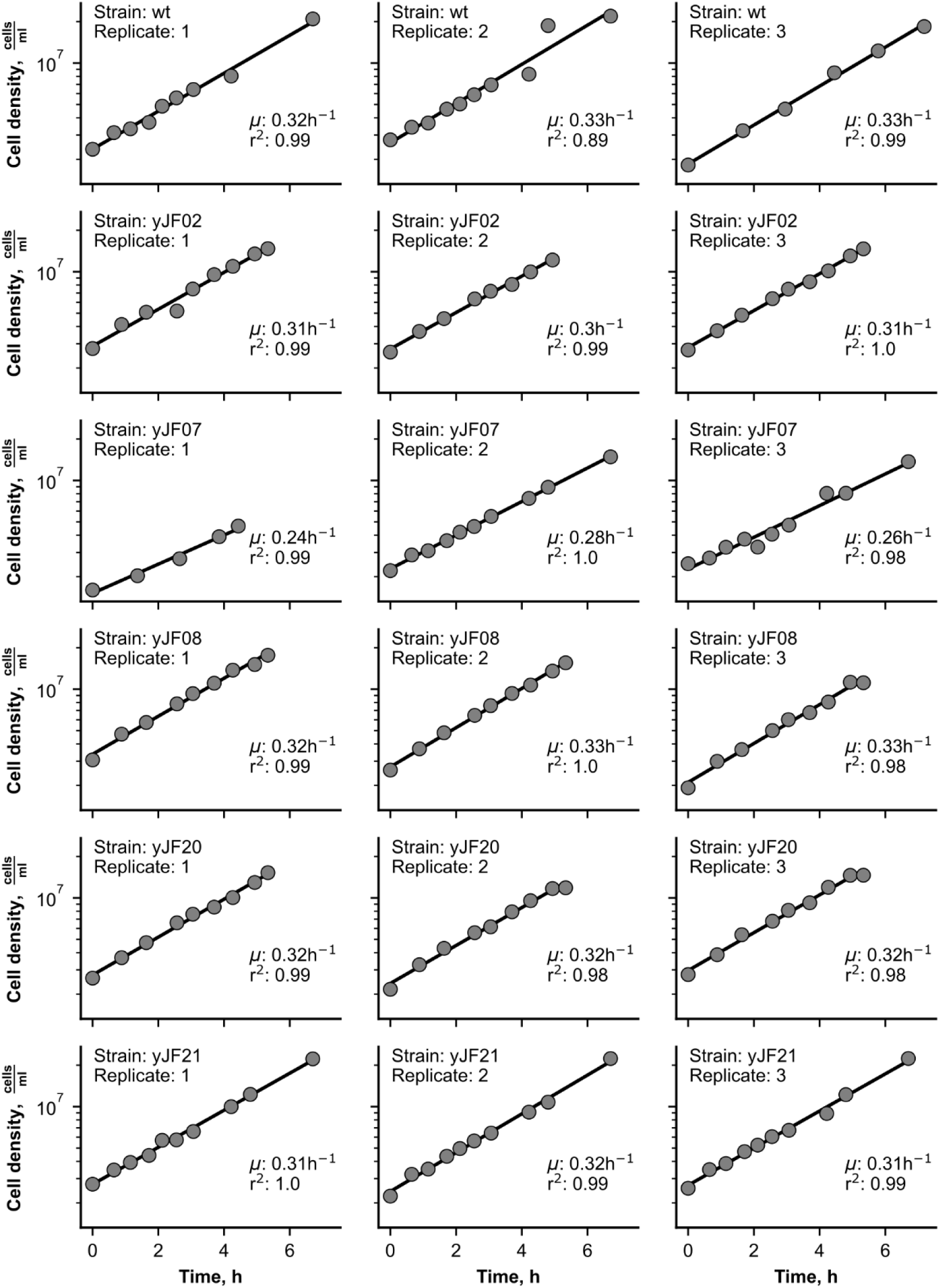
Determination of growth rates in each strain per experimental replicate. Solid lines indicate fits of a linear regression of the cell density measurements on a semi-logarithmic scale.

## References

1. Nishimura, K., Fukagawa, T., Takisawa, H., Kakimoto, T. & Kanemaki, M. An auxin-based degron system for the rapid depletion of proteins in nonplant cells. Nat Methods 6, 917–922 (2009).

2. Yesbolatova, A. et al. The auxin-inducible degron 2 technology provides sharp degradation control in yeast, mammalian cells, and mice. Nat Commun 11, (2020).

3. Kubota, T., Nishimura, K., Kanemaki, M. T. & Donaldson, A. D. The Elg1 Replication Factor C-like Complex Functions in PCNA Unloading during DNA Replication. Mol Cell 50, 273–280 (2013).

4. Holland, A. J., Fachinetti, D., Han, J. S. & Cleveland, D. W. Inducible, reversible system for the rapid and complete degradation of proteins in mammalian cells. Proc Natl Acad Sci U S A 109, (2012).

5. Maric, M., Maculins, T., De Piccoli, G. & Labib, K. Cdc48 and a ubiquitin ligase drive disassembly of the CMG helicase at the end of DNA replication. Science (1979) 346, (2014).

6. Nora, E. P. et al. Targeted Degradation of CTCF Decouples Local Insulation of Chromosome Domains from Genomic Compartmentalization. Cell 169, (2017).

7. Gibcus, J. H. et al. A pathway for mitotic chromosome formation. Science (1979) 359, (2018).

8. Assmus, H. E., Herwig, R., Cho, K. H. & Wolkenhauer, O. Dynamics of biological systems: Role of systems biology in medical research. Expert Review of Molecular Diagnostics vol. 6 891–902 Preprint at 10.1586/14737159.6.6.891 (2006).

9. Papagiannakis, A., Niebel, B., Wit, E. C. & Heinemann, M. Autonomous Metabolic Oscillations Robustly Gate the Early and Late Cell Cycle. Mol Cell 65, 285–295 (2017).

10. Papagiannakis, A., De Jonge, J. J., Zhang, Z. & Heinemann, M. Quantitative characterization of the auxin-inducible degron: A guide for dynamic protein depletion in single yeast cells. Sci Rep 7, (2017).

11. Takhaveev, V. et al. Temporal segregation of biosynthetic processes is responsible for metabolic oscillations during the budding yeast cell cycle. Nat Metab 5, 294–313 (2023).

12. Valenti, R. et al. A proteome-wide yeast degron collection for the dynamic study of protein function. Journal of Cell Biology 224, (2025).

13. Lehmayer, L., Bernauer, L. & Emmerstorfer-Augustin, A. Applying the auxin-based degron system for the inducible, reversible and complete protein degradation in Komagataella phaffii. iScience 25, (2022).

14. Tan, X. et al. Mechanism of auxin perception by the TIR1 ubiquitin ligase. Nature 446, 640–645 (2007).

15. Cao, S. Lessons from natural molecular glue degraders. Biochemical Society Transactions vol. 52 1191–1197 Preprint at 10.1042/BST20230836 (2024).

16. Yesbolatova, A., Natsume, T., Hayashi, K. ichiro & Kanemaki, M. T. Generation of conditional auxin-inducible degron (AID) cells and tight control of degron-fused proteins using the degradation inhibitor auxinole. Methods 164–165, (2019).

17. Mendoza-Ochoa, G. I. et al. A fast and tuneable auxin-inducible degron for depletion of target proteins in budding yeast. Yeast 36, 75–81 (2019).

18. Tanaka, S., Miyazawa-Onami, M., Iida, T. & Araki, H. iAID: An improved auxin-inducible degron system for the construction of a ‘tight’ conditional mutant in the budding yeast Saccharomyces cerevisiae. Yeast 32, (2015).

19. Watson, A. T., Hassell-Hart, S., Spencer, J. & Carr, A. M. Rice (Oryza sativa) tir1 and 5′adamantyl-iaa significantly improve the auxin-inducible degron system in schizosaccharomyces pombe. Genes (Basel) 12, (2021).

20. Natsume, T., Kiyomitsu, T., Saga, Y. & Kanemaki, M. T. Rapid Protein Depletion in Human Cells by Auxin-Inducible Degron Tagging with Short Homology Donors. Cell Rep 15, 210–218 (2016).

21. Nishimura, K. et al. A super-sensitive auxin-inducible degron system with an engineered auxin-TIR1 pair. Nucleic Acids Res 48, (2020).

22. Morawska, M. & Ulrich, H. D. An expanded tool kit for the auxin-inducible degron system in budding yeast. Yeast 30, 341–351 (2013).

23. Lee, M. E., DeLoache, W. C., Cervantes, B. & Dueber, J. E. A Highly Characterized Yeast Toolkit for Modular, Multipart Assembly. ACS Synth Biol 4, 975–986 (2015).

24. Santangelot, G. M. & Tornowt, J. Efficient Transcription of the Glycolytic Gene ADHI and Three Translational Component Genes Requires the GCRI Product, Which Can Act Through TUF/GRF/RAP Binding Sites. MOLECULAR AND CELLULAR BIOLOGY (1990).

25. Ho, B., Baryshnikova, A. & Brown, G. W. Unification of Protein Abundance Datasets Yields a Quantitative Saccharomyces cerevisiae Proteome. Cell Syst 6, (2018).

26. Zacharias, D. A., Violin, J. D., Newton, A. C. & Tsien, R. Y. Partitioning of lipid-modified monomeric GFPs into membrane microdomains of live cells. Science (1979) 296, (2002).

27. Lackner, D. H. et al. A Network of Multiple Regulatory Layers Shapes Gene Expression in Fission Yeast. Mol Cell 26, 145–155 (2007).

28. Wakil, S. J., Stoops, J. K. & Joshi, V. C. Fatty acid synthesis and its regulation. Annu Rev Biochem 52, 537–579 (1983).

29. Tehlivets, O., Scheuringer, K. & Kohlwein, S. D. Fatty acid synthesis and elongation in yeast. Biochimica et Biophysica Acta - Molecular and Cell Biology of Lipids vol. 1771 255–270 Preprint at 10.1016/j.bbalip.2006.07.004 (2007).

30. Schamhart, D. H. J., Ten Berge, A. M. A. & Van De Poll, K. W. Isolation of a Catabolite Repression Mutant of Yeast as a Revertant of a Strain That Is Maltose Negative in the Respiratory-Deficient State. JOURNAL OF BACTERIOLOGY vol. 121 https://journals.asm.org/journal/jb (1975).

31. Cherrick, H., Fugit, D. & Mowshowitzi, D. B. Pleiotropic Properties of a Yeast Mutant Insensitive to Catabolite Repression. (1980).

32. Lemontti, J. F., Fugit, D. R. & Mackayz, V. L. Pleiotropic Mutations at the TUP1 Locus That Affect the Expression of Mating-Type-Dependent Functions in Saccharomyces Cerevisiae. (1980).

33. Fu, S. F. et al. Indole-3-acetic acid: A widespread physiological code in interactions of fungi with other organisms. Plant Signaling and Behavior vol. 10 Preprint at 10.1080/15592324.2015.1048052 (2015).

34. Nicastro, R. et al. Indole-3-acetic acid is a physiological inhibitor of TORC1 in yeast. PLoS Genet 17, (2021).

35. Wang, X., Xu, H., Ha, S. W., Ju, D. & Xie, Y. Proteasomal degradation of Rpn4 in Saccharomyces cerevisiae is critical for cell viability under stressed conditions. Genetics 184, 335–342 (2010).

36. Canelas, A. B. et al. Integrated multilaboratory systems biology reveals differences in protein metabolism between two reference yeast strains. Nat Commun 1, (2010).

37. Pedregosa, F. et al. Scikit-learn: Machine learning in Python. Journal of Machine Learning Research 12, (2011).

38. Virtanen, P. et al. SciPy 1.0: fundamental algorithms for scientific computing in Python. Nat Methods 17, (2020).

39. Terpilowski, M. scikit-posthocs: Pairwise multiple comparison tests in Python. J Open Source Softw 4, (2019).

40. Sambrook, J. & Russell, D. W. The Inoue Method for Preparation and Transformation of Competent E. Coli: ‘Ultra-Competent’ Cells.

41. Sambrook, J. & Russell, D. W. Molecular Cloning: A Laboratory Manual. Third Edition. United States: Cold Spring Harbor Laboratory Press. (2001).

42. Novarina, D., Koutsoumpa, A. & Milias-Argeitis, A. A user-friendly and streamlined protocol for CRISPR/Cas9 genome editing in budding yeast. STAR Protoc 3, (2022).

43. Fan, L. CODON OPTIMIZATION. (2020).

44. Gietz, R. D. & Schiestl, R. H. Frozen competent yeast cells that can be transformed with high efficiency using the LiAc/SS carrier DNA/PEG method. Nat Protoc 2, (2007).

45. Heigwer, F., Kerr, G. & Boutros, M. E-CRISP: Fast CRISPR target site identification. Nature Methods vol. 11 Preprint at 10.1038/nmeth.2812 (2014).

46. Concordet, J. P. & Haeussler, M. CRISPOR: Intuitive guide selection for CRISPR/Cas9 genome editing experiments and screens. Nucleic Acids Res 46, (2018).

47. Castillo-Hair, S. M. et al. FlowCal: A User-Friendly, Open Source Software Tool for Automatically Converting Flow Cytometry Data from Arbitrary to Calibrated Units. ACS Synth Biol 5, (2016).

48. Stern, A. D., Rahman, A. H. & Birtwistle, M. R. Cell size assays for mass cytometry. Cytometry Part A 91, (2017).

49. Byrd, R. H., Schnabel, R. B. & Shultz, G. A. Approximate solution of the trust region problem by minimization over two-dimensional subspaces. Math Program 40, (1988).

50. Harris, C. R. et al. Array programming with NumPy. Nature vol. 585 Preprint at 10.1038/s41586-020-2649-2 (2020).

51. Koutsoumpa, A. & Milias-Argeitis, A. Rapid and reversible regulation of cell cycle progression in budding yeast using optogenetics. (2024) doi:10.1101/2024.09.21.614242.

